# Activation of the ChvG-ChvI pathway promotes multiple survival strategies during cell wall stress in *Agrobacterium tumefaciens*

**DOI:** 10.1101/2024.12.10.627833

**Authors:** Jacob M. Bouchier, Emily Knebel, Jennifer Amstutz, Gabriel Torrens, Gustavo Santiago-Collazo, Carli McCurry, Alexandra J. Weisberg, Felipe Cava, Pamela J.B. Brown

## Abstract

*Agrobacterium tumefaciens* shifts from a free-living soil bacterium to a plant-invading state upon encountering the plant root microenvironment. The acid-induced two-component sensor system ChvG-ChvI drives this shift and triggers a complex transcriptional program that promotes host invasion and survival against host immune defenses. Remarkably, ChvG-ChvI is also activated under cell wall stress conditions suggesting that the transcriptional response may have a broader function. Here, we find that blocking cell wall synthesis either genetically or chemically leads to ChvG-ChvI activation. Mutations in key cell wall synthesis or outer membrane proteins, such as PBP1a, FtsW, and AopA1, suppress ChvG-ChvI activation suggesting that providing structural integrity is a primary function of the ChvG-ChvI regulon. Here, we investigated regulon components for this function. First, the exopolysaccharide succinoglycan confers tolerance to multiple β-lactam antibiotics targeting different enzymes by forming a protective barrier around the cells. Next, a Class D β-lactamase is expressed which may contribute to the high level of β-lactam resistance in *A. tumefaciens*. Finally, outer membrane remodeling compensates for the accumulation of cell wall damage by providing structural integrity. Overall, we expand our understanding of mechanisms driving ChvG-ChvI activation and β-lactam resistance in a bacterial plant pathogen.

**Significance Statements.:**

- Activation of the ChvG-ChvI two component system promotes survival when the bacterial cell walls are damaged by a variety of genetic or chemical approaches
- The ChvG-ChvI dependent production of the exopolysaccharide succinoglycan, β-lactamase Cbl activity, and outer membrane proteome remodeling all contribute to survival in the presence of β-lactam antibiotics
- Improved understanding of bacterial stress responses that promote antibiotic tolerance and resistance has the potential to inform development of novel drug targets

## Introduction

Central to their survival, bacteria exhibit remarkable phenotypic plasticity, employing a myriad of survival strategies in response to both biotic and abiotic stressors in their environment. One strategy they employ for environmental sensing is the two-component system (TCS). TCSs are robust signal/response mechanisms that transduce external stimuli into transcriptional responses (Capra and Laub, 2012). One well-described TCS conserved in many, but not all the Alphaproteobacteria is the sensor kinase and response regulator pair ChvG-ChvI (Charles and Nester, 1993).

ChvG-ChvI was first described in the plant symbiont *Sinorhizobium meliloti* (ExoS-ChvI) and the plant pathogen *Agrobacterium tumefaciens*. Initially, this system was found to be activated in acidic conditions, leading to a signal cascade that results in differential expression of hundreds of genes (Yuan *et al*., 2008). ChvG-ChvI is repressed by the accessory protein ExoR. In acidic conditions, ExoR is proteolyzed and disassociates from ChvG, enabling activation (Lu *et al*., 2012; Heckel *et al*., 2014). ChvG-ChvI is also required for plant association, likely playing a crucial role in the transition to a host- invading state. This host-invading function with a link to pH-driven activation of ChvG-ChvI is conserved in the symbiotic and pathogenic species of the Hyphomicrobiales and is well described in Greenwich et al., 2023. However, recent work in *Caulobacter crescentus* and *A. tumefaciens* expands the conditions which activate ChvG-ChvI to include osmotic and cell wall stress (Quintero-Yanes *et al*., 2022; Williams *et al*., 2022).

Depletion of an essential cell wall synthase, penicillin-binding protein 1a (PBP1a), in *A. tumefaciens* leads to changes in the transcriptional profile which align with the known ChvG-ChvI regulon (Williams *et al*., 2022). PBP1a is a bifunctional class A penicillin-binding protein that drives polar elongation in *A. tumefaciens* using glycosyltransferase and transpeptidase activities to build the bacterial cell wall, which is made of a meshwork of crosslinked peptidoglycan (PG) strands (Williams *et al*., 2021). The bacterial cell wall is essential for maintaining cell shape and resistance to osmotically unfavorable environments. Proper cell wall synthesis requires a high degree of coordination amidst many PG biosynthesis and remodeling enzymes. While there is some flexibility in withstanding a small degree of cell wall damage, larger defects are typically lethal (Huang *et al*., 2008). Thus, we hypothesize that cell wall damage, rather than loss of any one enzyme activity, is likely to activate the ChvG-ChvI pathway.

Indeed, Δ*chvG* and Δ*chvI* mutants are hypersensitive to β-lactam antibiotics suggesting that activation of this two-component system provides protection when cell walls are compromised (Williams *et al*., 2022). Our study aims to better understand the activation of ChvG-ChvI during cell wall stress conditions in *A. tumefaciens*. Further, we sought to understand how the ChvG-ChvI response is protective against cell wall stress.

This study builds upon previous research by showing that inhibition of cell wall synthases involved in both polar elongation and cell division triggers the activation of ChvG-ChvI. Activation of ChvG-ChvI results in many physiological changes, including succinoglycan production. Here we show that succinoglycan acts as a protective barrier against stresses incurred by antibiotic-driven inhibition of cell wall synthesis. We show that succinoglycan provides protection against the β-lactam antibiotics. We characterize the contributions of a putative ChvG-ChvI regulated class D β-lactamase that modestly contributes β-lactam to resistance. Finally, we find that ChvG-ChvI-mediated modulation of the outer membrane proteome likely promotes structural integrity when the cell wall is damaged. Overall, our study expands the knowledge of conditions that activate ChvG-ChvI and the benefits conferred to *A. tumefaciens* cells under an array of cell wall stress conditions.

## Results

### Treatment with ampicillin and cefsulodin activates the ChvG-ChvI two-component system through distinct mechanisms

Deletion mutants of *chvG* and *chvI* in *Agrobacterium tumefaciens* are hypersensitive to the β-lactam cefsulodin, which inhibits PG synthesis. Cefsulodin treatment of wild-type cells leads to short, round cells and cell spreading from succinoglycan overproduction, consistent with decreased PBP1a activity (Williams *et al*., 2022). To better understand this hypersensitivity, we generated a cefsulodin suppressor mutant by adaptive evolution in our Δ*chvI* background (Δ*chvI*^cef^). This Δ*chvI*^cef^ strain was subsequently confirmed to grow as well as Wild type in 25 µg/ml of cefsulodin (Figure 1A).

**Figure 1.**
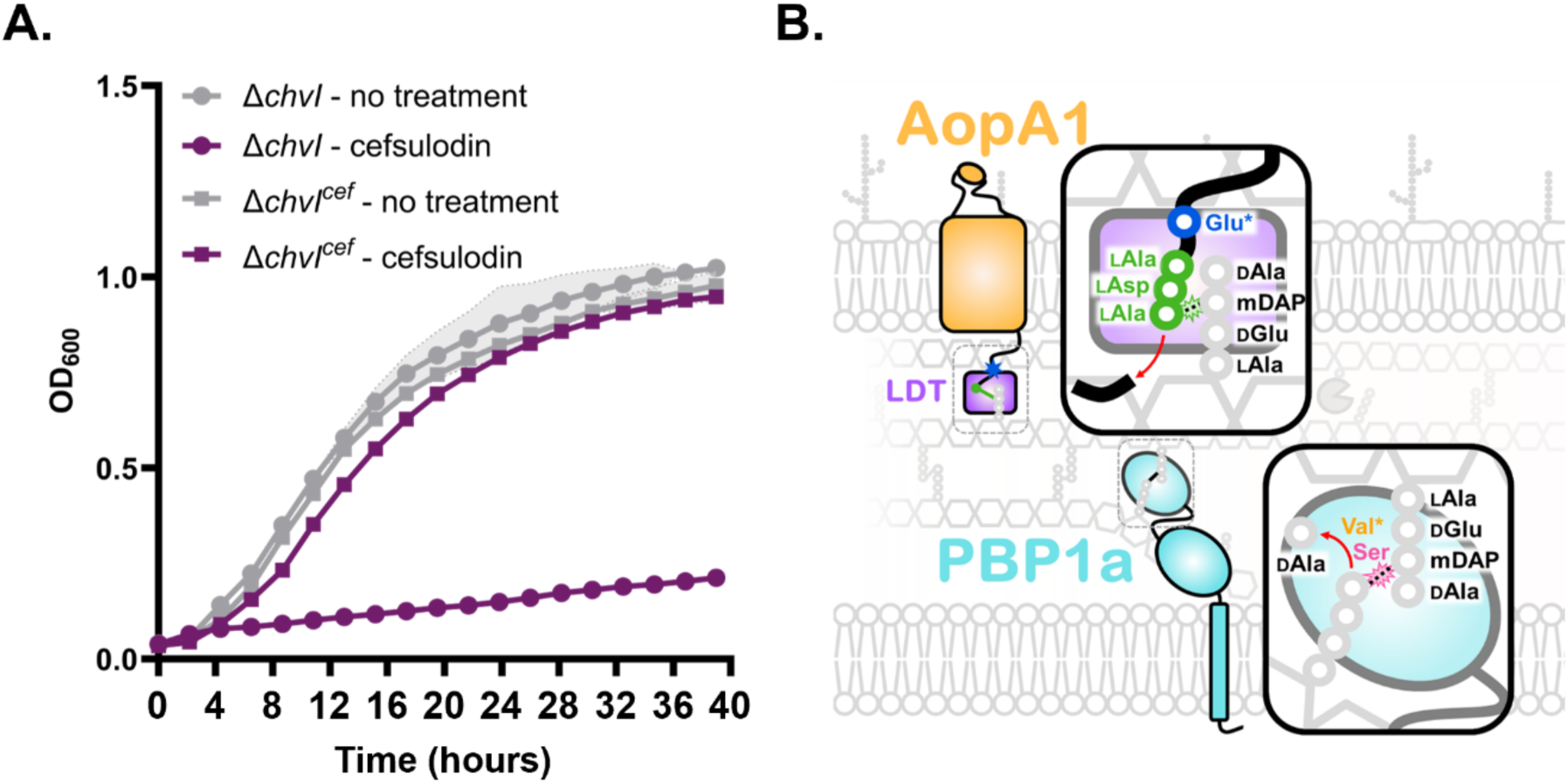
Cefsulodin-adapted Δ*chvI* strain (Δ*chvI^cef^*) possess two missense mutations in essential genes *aopA1* and *mrcA*. A) Growth curve of Δ*chvI* (circles) and *chvI^c^*^ef^ (squares) in ATGN media containing either 25 µg/ml of cefsulodin or no antibiotic measured by OD_600_ over 40 hours. Data shown represents the average of 3 biological replicates, with shaded regions indicating standard deviation. B) Schematic of the point mutations identified from variant analysis of the Δ*chvI^c^*^ef^ strain genome compared to wildtype (WT). *aopA1* (Atu1020) is predicted to encode an essential porin. The Glu32Gly mutation in Δ*chvI^cef^* (WT glutamate shown in blue) is 7 amino acids downstream of the ADA sequence (green residues), which is the dependent site for ld- transpeptidase-driven crosslinking of β-barrel proteins to the peptidoglycan. *mrcA* (Atu1341) encodes the essential bifunctional penicillin binding protein 1a, where there is a Val659Met mutation in Δ*chvI^cef^*. The wildtype valine residue (shown in orange) is sterically near the catalytic active site (represented by the pink Ser) of the transpeptidase domain in the folded structure. Green and pink starbursts indicate the crosslinking mediated by ld-transpeptidases and PBP1a, respectively.

Whole genome sequencing and subsequent analysis identified deletions that are present in our lab strains relative to the published *Agrobacterium tumefaciens* C58 reference genome (NCBI: GCF_000092025.1), confirmed the presence of the *chvI* deletion, and identified other deletions that arose during passaging. However, none of the deletions were unique to our Δ*chvI*^cef^ strains (Supplemental Tables 1-2). Single nucleotide polymorphism (SNP) variant analysis identified mutations that arose during passaging of Wild type or Δ*chvI* without the antibiotic, as well as one synonymous and two missense mutations unique to Δ*chvI*^cef^ (Supplemental Table 3). The first missense mutation in Δ*chvI*^cef^ was in the essential outer membrane porin protein AopA1 (encoded by *atu1020*), which has been auto-annotated as important for the passive diffusion of small molecules (Curtis and Brun, 2014; Paysan-Lafosse *et al*., 2023). The mutation resulted in a change in the coding sequence from a glutamate to a glycine at residue 32 (E32G) in the disordered, periplasmic-exposed region of the protein (Figure 1B; Supplemental Figure 1). Notably, this residue is seven amino acids from the ADA sequence required for LD-transpeptidase (LDT)-mediated cross-linking to PG (Godessart *et al*., 2020; Sandoz *et al*., 2021). The second mutation in Δ*chvI*^cef^ resulted in a change in the coding sequence of PBP1a (encoded as *mrcA*) at residue 659, topologically close to the active site of the transpeptidase domain (Figure 1B; Supplemental Figure 1). The change was from a valine to a bulkier methionine (V659M), suggesting that a local change in charge or steric hindrance may interfere with the binding of cefsulodin. AlphaFold2 structure predictions and electrostatic modeling of the transpeptidase domain of PBP1a^V659M^ support this hypothesis (Supplemental Figure 2). Replacement of wildtype *mrcA* with a variant that encodes PBP1a^V659M^ at the native locus abolishes the overproduction of succinoglycan during treatment with 25 µg/mL cefsulodin and is sufficient to restore growth to Δ*chvG* and Δ*chvI* in defined media containing 25 µg/mL cefsulodin (Figure 2, Supplemental Figure 3). These observations suggest that cefsulodin-mediated inactivation of PBP1a is sufficient to activate ChvG- ChvI.

**Figure 2.**
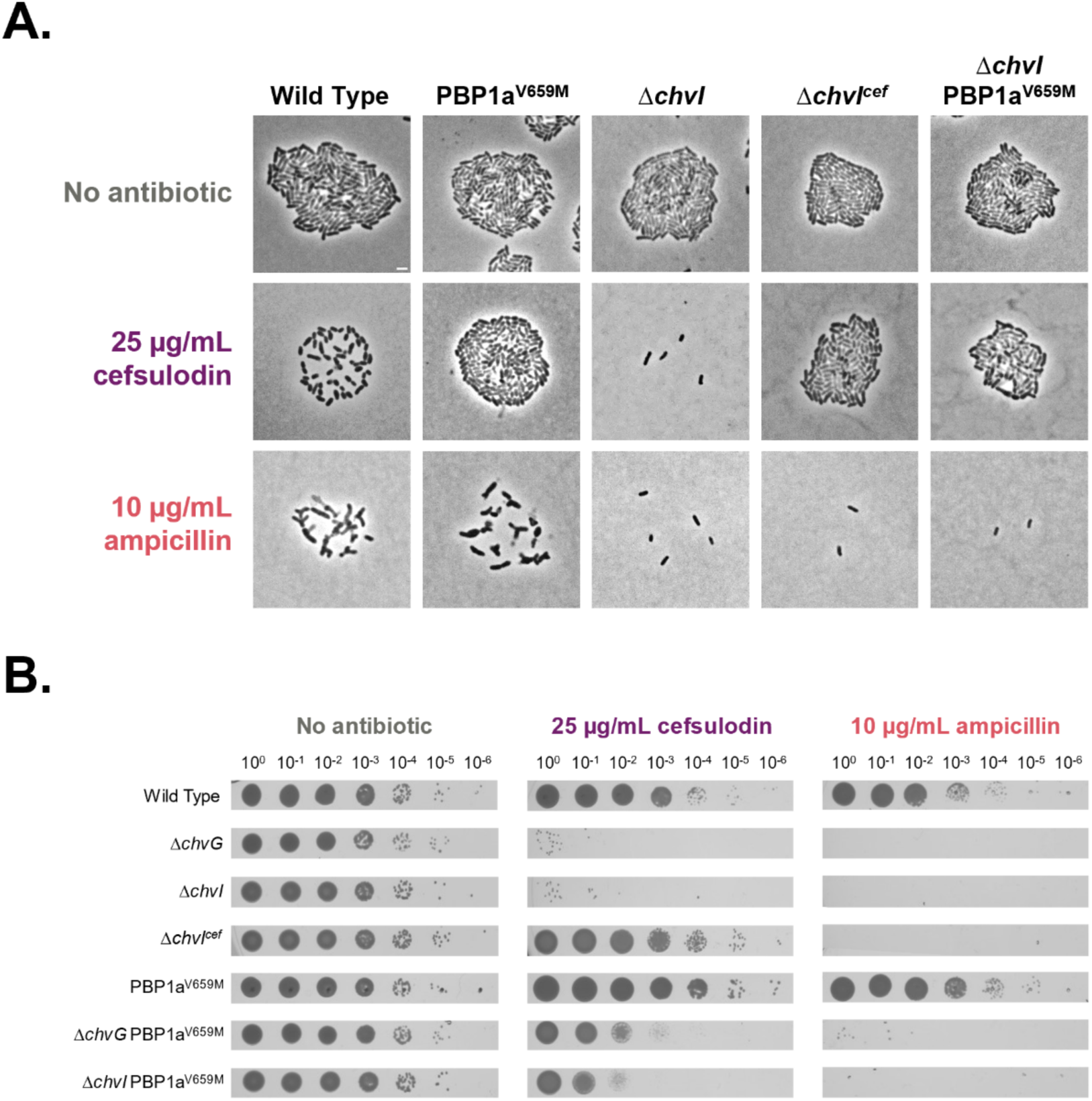
PBP1a^V659M^ confers cefsulodin-specific protection. A) Growth after 16 hours on a 1.5% ATGN agarose pad with no antibiotic, 25 µg/ml cefsulodin, or 10 μg/mL ampicillin. Spreading microcolonies are indicative of succinoglycan production following treatment with antibiotics that activate ChvGI. Scale bar represents 2 µm. B) Cell viability for each strain on ATGN plates containing no antibiotic, 25 µg/ml cefsulodin, or 10μg/mL ampicillin is assessed by spotting serial dilutions incubated for 3 days.

Next, we sought to determine if the PBP1a^V659M^ allele would confer resistance to other β-lactam antibiotics. As seen with cefsulodin treatment, we observed cell spreading indicative of ChvG-ChvI activation during overnight treatment with 10 µg/mL ampicillin in wild-type cells and hypersensitivity in Δ*chvG* and Δ*chvI* (Figure 2A). Ampicillin treatment resulted in cellular elongation and branching of wild-type cells, a morphology distinct from the short, round cells observed during cefsulodin treatment (Figure 2A). The branched morphologies are similar to cells previously observed during inhibition of septal peptidoglycan (sPG) synthesis in *A. tumefaciens* (Howell *et al*., 2019; Williams *et al*., 2021; Figueroa-Cuilan *et al*., 2022). The cefsulodin-suppressing PBP1a^V659M^ allele was not sufficient to reduce overproduction of succinoglycan during treatment with ampicillin (Figure 2A). Further, *chvG* and *chvI* mutants containing the PBP1a^V659M^ allele remained hypersensitive to ampicillin (Figure 2, Supplemental Figure 3). These observations led us to suspect that activation of ChvG-ChvI may occur during a variety of stressors that impact cell wall integrity, rather than inhibition of PBP1a specifically.

### The ChvG-ChvI two-component system is transcriptionally activated by the presence of cell wall targeting antibiotics

To assess ChvG-ChvI activity, we constructed a P*_chvGI_* -driven *lacZ* reporter plasmid to quantify ChvG-ChvI promoter activity by measuring β-galactosidase activity. We validated that our P*_chvGI_* reporter functions as expected using two constitutively active ChvG-ChvI strains, *ΔexoR* and phosphomimetic *chvI*^D52E^, and three ChvG-ChvI deactivating strains, *ΔchvG, #x0394;chvI,* and phosphoablative *chvI*^D52N^. We hypothesized that constitutive ChvG-ChvI upregulation would enable growth in the presence of cell wall targeting antibiotics but the phosphoablative mutant would be hypersensitive. Indeed, the *chvI*^D52N^ strain was hypersensitive to both antibiotics (Figure 3A), suggesting that phosphoablation of ChvI is sufficient to block ChvG-ChvI-mediated protection against these antibiotics. Further, *chvI*^D52E^ and *ΔexoR* strains both exhibited increased production of succinoglycan as evidenced by cell spreading, while *ΔchvG* and *chvI*^D52N^ do not spread (Figure 3B). As expected, the absence of ExoR or the presence of the phosphomimic ChvI^D52E^ leads to increased P*_chvGI_* activity and ChvI^D52N^ does not (Figure 3C; Supplemental Table 4).

**Figure 3.**
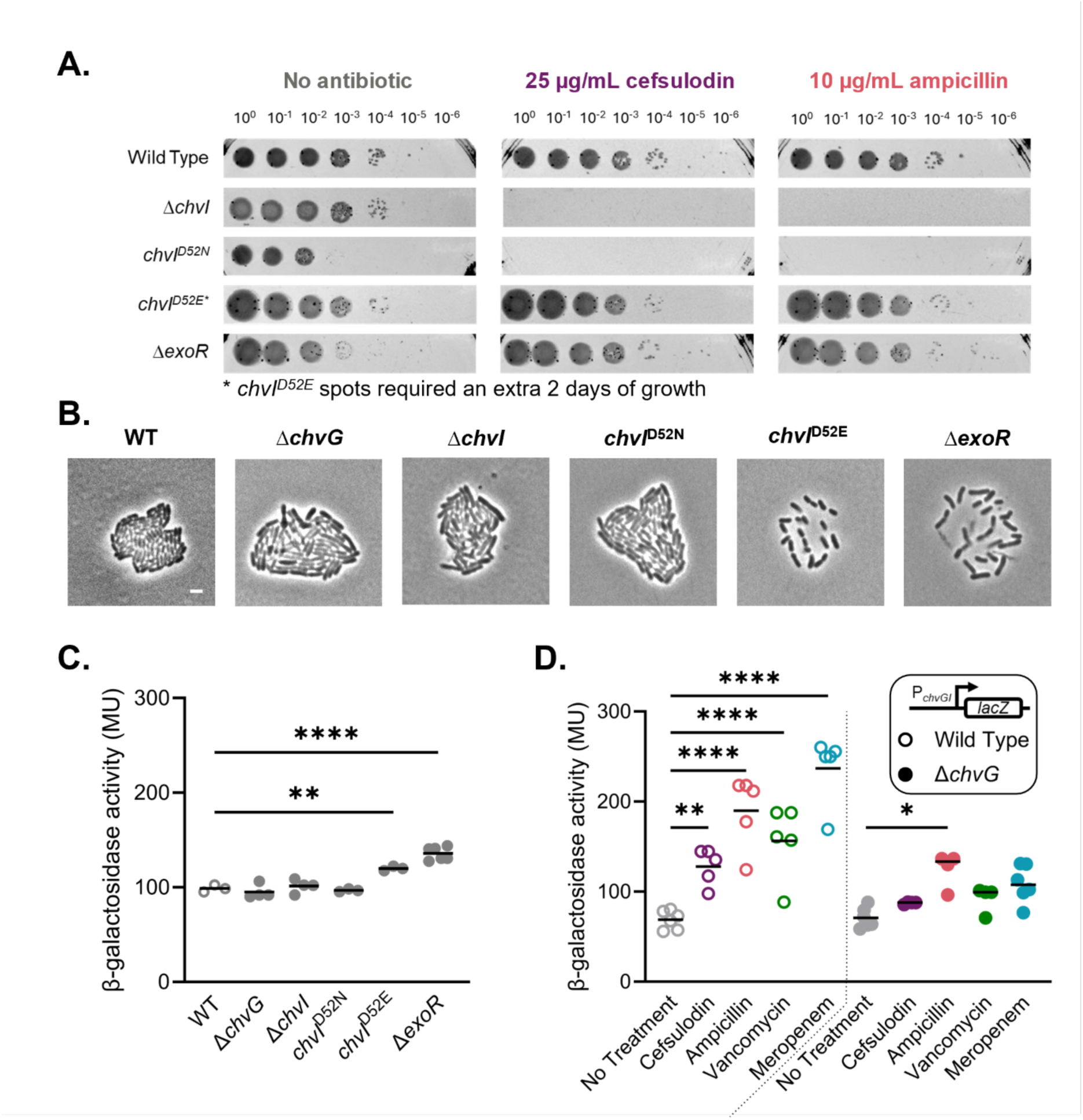
*chvG-chvI* promoter expression is upregulated in constitutively active mutants and in the presence of cell wall targeting antibiotics, which restores beta lactam resistance. A) Cell viability for each strain on ATGN plates containing no antibiotic, 25 µg/ml cefsulodin, or 10 μg/mL ampicillin is assessed by spotting serial dilutions for 3 days (*except for *chvI^D52E^*, which incubated for 5 days due to slower growth). B) Growth after 16 hours on a 1.25% ATGN agarose pad. ChvG-I activation results in the production of less compact microcolonies. Scale bar represents 2 µm. C) β-galactosidase activity in Miller Units (MU) as a reporter of *chvGI* promoter activity in ChvG-ChvI activating or inactivating strains grown from exponential phase for 6 hours in ATGN. Chromosomal *chvI* alleles encoding the phosphomimetic D52E and phosphoablative D52N variants are expressed from the native *chvI* locus. Significance compares each mean and was calculated with an ANOVA and Tukey’s post hoc test. *, p < 0.05; **, p < 0.005; ***, p < 0.0005. Nonsignificant comparison backets are not shown. D) β-galactosidase activity as a reporter of *chvGI* promoter activity in antibiotic conditions. Exponential phase WT (open circles) and Δ*chvG* (closed circles) incubated with 20 µg/ml of each antibiotic for 6 hours. Significance compares each treatment to the no treatment control and was calculated with an ANOVA and Tukey’s post hoc test. ****, p < 0.00005. Nonsignificant comparison bars were not shown.

To determine if P*_chvGI_* activity is increased when PG synthesis is impaired, P*_chvGI_* activity was measured with our β-galactosidase reporter after a 6-hour treatment with a variety of cell wall targeting antibiotics. Consistent with the observation of wild-type cell spreading when treated with cefsulodin and ampicillin (Figure 2A) and hypersensitivity of *ΔchvG* and *ΔchvI* strains to these drugs (Figure 2B; Williams et al., 2022), P*_chvGI_* activity is increased when cells are treated with these two β-lactam antibiotics (Figure 3D, Supplemental Table 5). Increased P*_chvGI_* activity was also detected during treatment with vancomycin, a glycopeptide that binds the terminal D-alanines on the muropeptide stem, preventing cell wall synthesis (Figure 3D, Supplemental Table 5). This observation is consistent with our demonstration that PBP1a activity alone is not responsible for ChvG-ChvI activation, but rather a general impairment of cell wall synthesis.

Additionally, we tested the impact of meropenem, also a β-lactam, treatment on P*_chvGI_* activity (Figure 3D, Supplemental Figure 4 & 6A, Supplemental Tables 5-6). Treatment with a low concentration of meropenem (1.5 μg/ml) causes dramatic mid-cell swelling (Supplemental Figure 4D) and results in a decrease in LD-crosslinks in the PG (Williams *et al*., 2022) suggesting that at this concentration meropenem targets LD-transpeptidases (LDTs). Increased P*_chvGI_* activity was not detected with sublethal meropenem treatments for 6 hours (1.5 μg/ml) although this concentration is sufficient to significantly inhibit cell growth in wild-type cells (Supplemental Figure 4A). In our Δ*chvG* mutant, 1.5 µg/ml of meropenem completely inhibits growth, with a significant growth inhibition at 0.75 µg/ml, which mildly impacted growth in wild-type cells, affirming that Δ*chvG* mutants are hypersensitive to meropenem (Supplemental Figure 4A-B).

Remarkably, when meropenem is present at a high concentration (20 μg/ml) during 6 hours of incubation, significant P*_chvGI_* activity was detected despite this concentration well-exceeding the minimum inhibitory concentration of 3 µg/ml in Wild type (Figure 3C, Supplemental Figure 4A & C, Supplemental Tables 5-6). This elevated promoter activity at high meropenem concentrations is perhaps due to the inactivation of LDTs and other cell wall synthases including PBPs. Together, these observations suggest that growth inhibition is not sufficient for ChvG-ChvI activation but rather that perturbation of cell wall integrity through disruption of PG synthases involved in elongation or cell division (Figure 4A) may lead to ChvG-ChvI activation.

**Figure 4.**
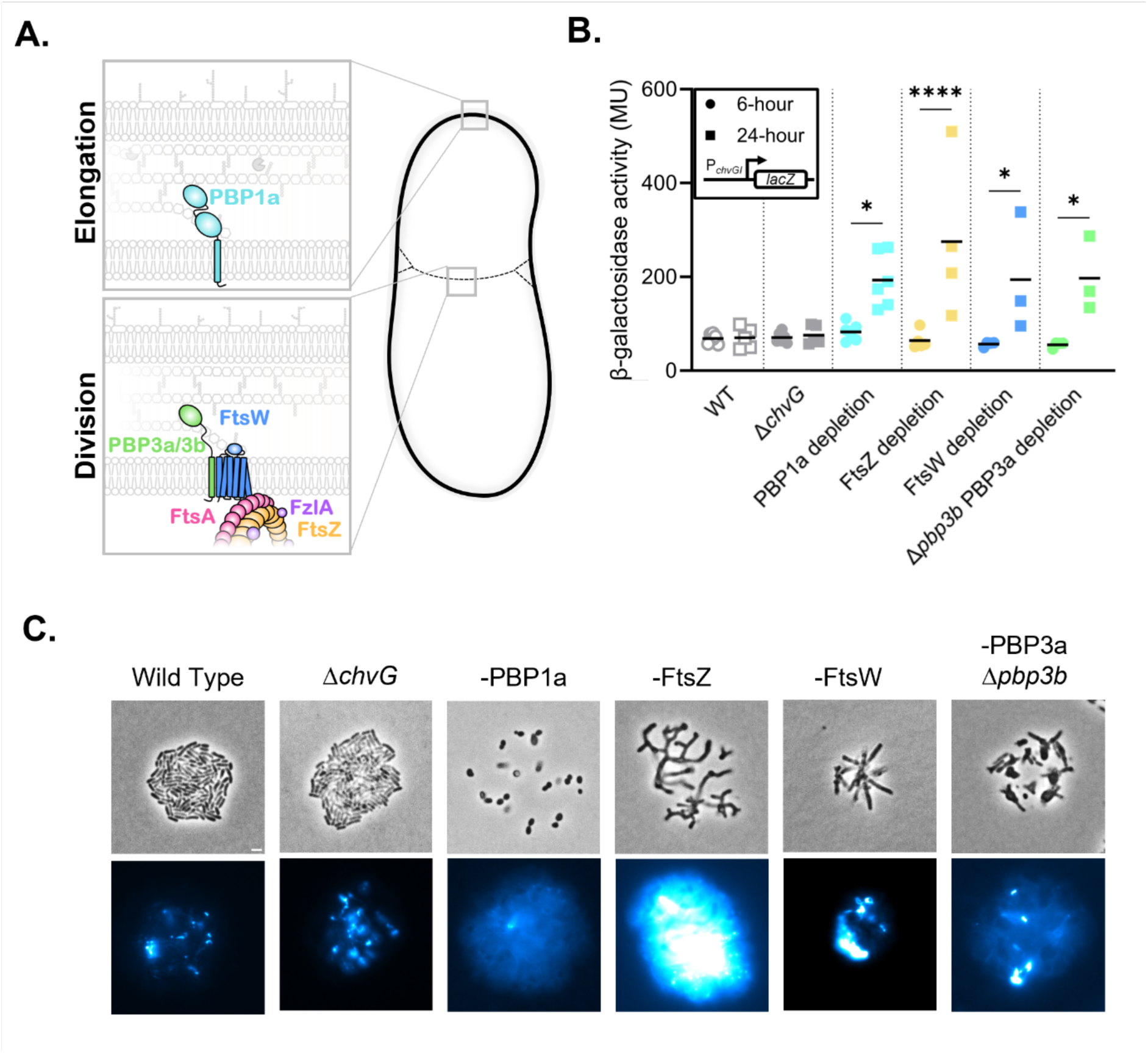
ChvG-ChvI is activated during depletion of essential elongation and division proteins. A) Schematic depicting subcellular locations of PG synthases and associated regulatory proteins. B) β-galactosidase activity in Miller Units (MU) as a reporter of *chvGI* promoter activity in each indicated strain measured after 6 hours and 24 hours of growth under depletion conditions. Bars represent the mean of the represented data. Significance compares 24-hour measurement to the corresponding 6- hour measurement and was calculated with an ANOVA and Tukey’s post hoc test. *, p < 0.05; **, p < 0.005; ***, p < 0.0005. Nonsignificant comparisons are not shown. C) Microcolony production after 24 hours of growth on 1.5% ATGN agarose pad containing 200 µM calcofluor white to stain succinoglycan. Inoculation to the pad occurred after 8 hours of growth in liquid ATGN without inducer for the gene depletions. Top micrographs are in phase and bottom were taken using a DAPI filter and false colored using the ImageJ LUT preset “cyan hot”. All DAPI fluorescence channels were normalized to the same values. Scale bar represents 2 µm.

### Inhibition of septal PG synthesis activates ChvG-ChvI in *A. tumefaciens*

The branching morphologies and spreading microcolonies observed during ampicillin treatment (Figure 2A) suggests that inhibition of sPG biosynthesis may activate the ChvG-ChvI pathway. sPG biosynthesis is mediated by a shape, elongation, division, and sporulation (SEDS) family glycosyltransferase (GTase) functionally coupled to a monofunctional penicillin binding protein (bPBP) transpeptidase (TPase), comprising the SEDS-bPBP complex (Meeske et al., 2016; Rohs et al., 2018; Taguchi et al., 2019). The septal PG machinery in *A. tumefaciens* is comprised of FtsW and PBP3, which exist predominantly at midcell and function independently of elongation machinery such as PBP1a at the pole (Figure 4A). To test sPG inhibition more directly, we took advantage of depletion strains known to cause sPG synthesis defects in *A. tumefaciens* using the β-galactosidase assay as a reporter of P*_chvGI_* activation and calcofluor white staining to detect succinoglycan production as an output for ChvG-ChvI activation. As a control, we confirmed increased P*_chvGI_* activity (Figure 4B) and succinoglycan production (Figure 4C) in the PBP1a depletion (-PBP1a). Depletion of FtsZ, the essential initiator of cell division, results in elongated, branching cells (Figure 4C; Howell *et al.,* 2019) presumably due to the inability to recruit the sPG machinery. After 24 hours of FtsZ depletion, P*_chvGI_* activation (Figure 4B, Supplemental Table 7) and succinoglycan production (Figure 4C) are readily detected, consistent with ChvG-ChvI activation. Next, we sought to determine if ChvG-ChvI is activated if the sPG synthases are deleted or depleted. The *A. tumefaciens* genome encodes two PBP3 orthologs, PBP3a and PBP3b, and these enzymes have some degree of functional redundancy as both *pbp3a* and *pbp3b* can be deleted with minimal impacts on cell viability (Williams *et al*., 2021). Consistent with the maintenance of cell wall integrity in these strains, the deletion of *pbp3a* or *pbp3b* does not lead to activation of P*_chvGI_* (Supplemental Figure 5, Supplemental Table 8). In contrast, depletion of *pbp3a* in the Δ*pbp3b* background (- PBP3ab) causes mid-cell swelling and ectopic pole formation (Fig 4C; Williams *et al*., 2021) indicative of a block in sPG synthesis which is sufficient to activate the ChvG-ChvI pathway (Figure 4B-C). Similarly, depletion of *ftsW* (-FtsW), results in mid-cell swelling and ectopic pole formation (Figure 4C; Howell et al 2019) and activates the ChvG-ChvI pathway (Figure 4B-C, Supplemental Table 7). Together, these data suggest that disruption of the SEDS-bPBP sPG synthesis complex of FtsW-PBP3ab is sufficient to activate the ChvG-ChvI pathway.

In both *Caulobacter crescentus* and in *Agrobacterium tumefaciens,* sPG synthesis is regulated by the FtsZ-localized protein FzlA (Goley et al., 2010; Lariviere et al., 2019). Depletion of FzlA (-FzlA) in *A. tumefaciens* causes sPG synthesis defects similar to both -FtsW and -PBP3ab (Figure 5C; Lariviere 2019). Importantly, the sPG synthesis defects of -FzlA can be rescued by expressing a hyperactive variant FtsW (FtsW^F137L^) (Figure 5A,C; Lariviere et al., 2019), consistent with a function for FzlA in FtsW-PBP3ab activation. Like the other depletions that impair sPG synthesis, after 48 hours of depletion, -FzlA exhibited increased P*_chvGI_* activity and a qualitative increase in succinoglycan pooling (Figure 5B-C, Supplemental Table 9). These data suggest that increased FtsW activity was sufficient to reduce ChvG-ChvI activation during depletion of FzlA in *A. tumefaciens*. Overall, disruption of PG synthesis via multiple mechanisms can lead to activation of the ChvG-ChvI pathway.

**Figure 5.**
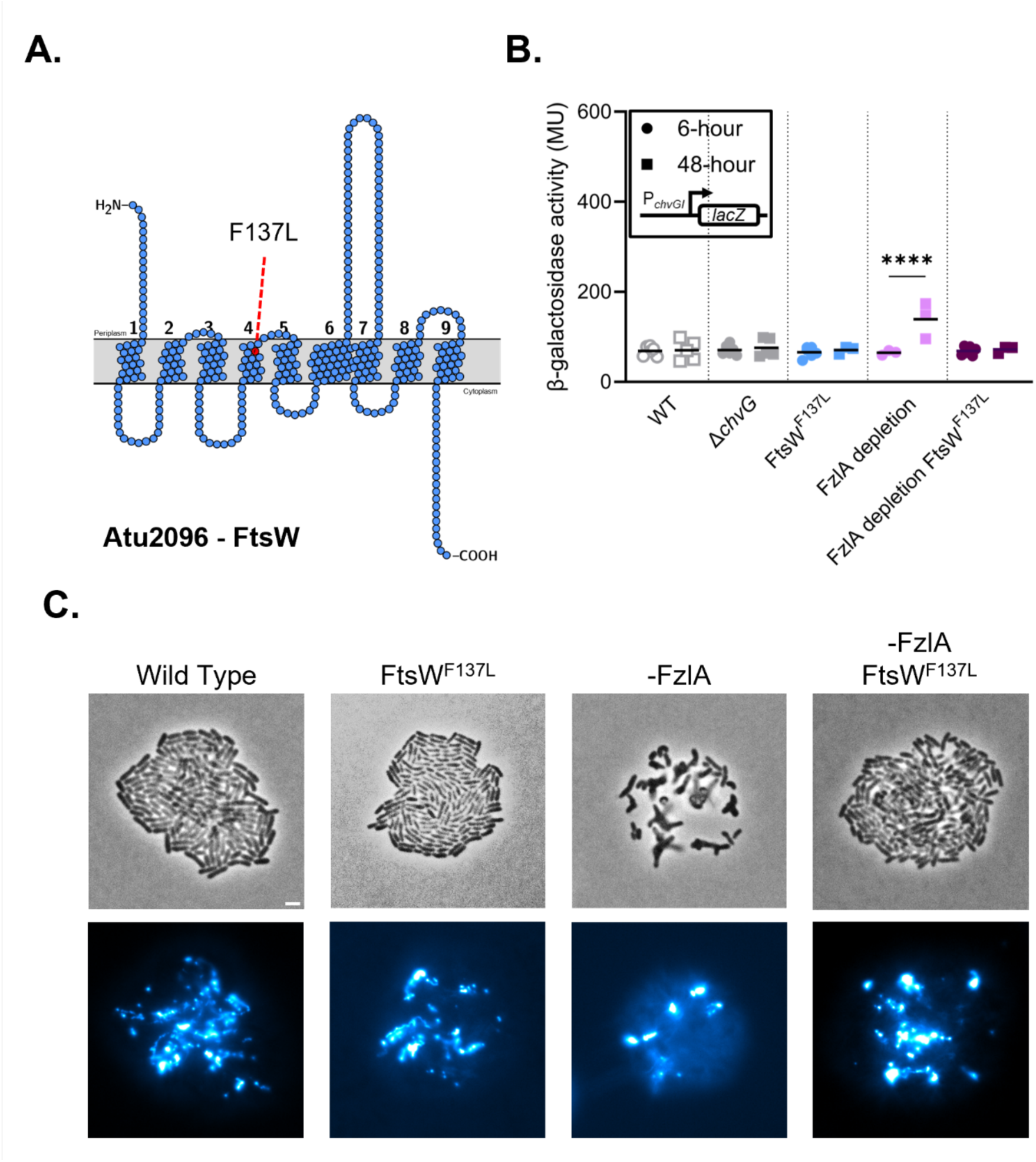
**ChvG-ChvI is activated during misregulation of sPG synthesis in *Agrobacterium tumefaciens.*** A) Schematic depicting topology maps of FtsW from *Agrobacterium tumefaciens* with red dashed lines indicating the location of the hyperactive mutant allele F137L. B) β-galactosidase activity in Miller Units (MU) of each strain measured at 6 and 48 hours of growth. Bars represent the mean of the represented data. Significance compares 24-hour measurement to the corresponding 6-hour measurement and was calculated with an ANOVA and Tukey’s post hoc test. *, p < 0.05; **, p < 0.005; ***, p < 0.0005. Nonsignificant comparisons are not shown C) Microcolony production after 24 hours of growth on 1.5% ATGN agarose pad containing 200 µM calcofluor white to stain succinoglycan. Inoculation to the pad occurred after 24 hours of growth in liquid ATGN without inducer for the gene depletions. Top micrographs are in phase and bottom were taken using a DAPI filter and false colored using the ImageJ LUT preset “cyan hot”. All DAPI fluorescence channels were normalized to the same values. Scale bar represents 2 µm.

### Succinoglycan and β-lactamase production confer protection against cell wall damage

The ChvG-ChvI regulon includes hundreds of genes with diverse functions including the repression of motility, increased succinoglycan production, and activation of type VI secretion (Greenwich *et al*., 2023). While ChvG-ChvI is clearly activated in an array of conditions that disrupt cell wall synthesis and compromise its integrity, it is less clear how transcriptional changes that occur during ChvG-ChvI activation provide an advantage. To explore how ChvG-ChvI activation may promote survival, we first considered a potential role for succinoglycan.

Succinoglycan biosynthesis genes are highly expressed under conditions that activate ChvG-ChvI (Figure 6A) and succinoglycan production is directly regulated by ChvG-ChvI activation (Ratib *et al*., 2018). Thus, we tested if succinoglycan production contributes to protection against external stressors, including the cell-wall targeting antibiotics. While cefsulodin, meropenem, and ampicillin are β-lactam antibiotics (Supplemental Figure 6A), treatment with sublethal concentrations of these drugs gives rise to distinct phenotypes in *A. tumefaciens* (Figure 2A and Figure Supplemental Figure 4D) suggesting they have different enzymatic targets. Thus, we hypothesized that if succinoglycan forms a protective barrier, it should confer tolerance to all three antibiotics.

**Figure 6.**
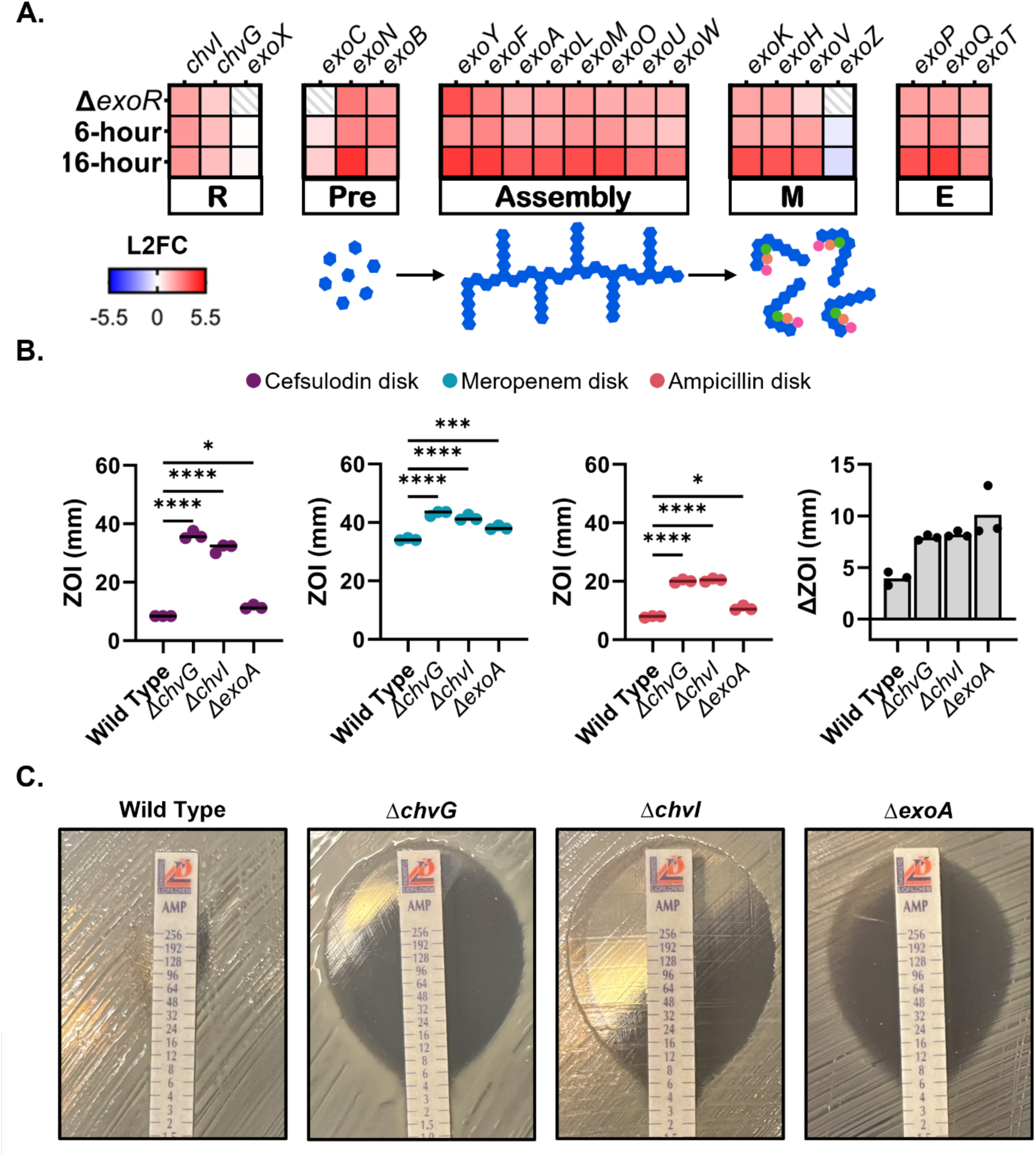
Contributions of ChvG-ChvI mediated succinoglycan to cefsulodin, meropenem, and ampicillin resistance. A) Log 2-fold change (L2FC) of genes involved in succinoglycan biosynthesis in three ChvG-ChvII activating conditions, deletion of *exoR* (Heckel et al., 2014) and depletion of PBP1a for 6- or 16-hours (Williams et al., 2022). Shading represents L2FC where downregulated genes are a gradient of blue and upregulated genes are a gradient of red. White represents no change and white with gray stripes represents L2FC values that were insignificant and not reported. Genes for precursor synthesis (Pre), polysaccharide assembly (Assembly), modification (M), and export (E) are shown, along with a simplified schematic of succinoglycan biosynthesis. B) Disk diffusion assays were conducted with cefsulodin (100 µg), meropenem (10 µg), ampicillin (10 µg) and ampicillin + sulbactam (10µg + 10µg) and the zones of inhibition (ZOI) for each strain are shown. The difference in ZOI between ampicillin + sulbactam and ampicillin (ΔZOI) is also shown. Comparison of means for cefsulodin and meropenem disks was performed with a one- way ANOVA followed by Tukey’s multiple comparisons test with a single pooled variance. Comparison of means for ampicillin and ampicillin + sulbactam disks was performed with a two-way ANOVA, followed by Šídák’s multiple comparisons test with a single pooled variance. ns, not significant; *, p<0.1; **, p<0.01; ***, p<0.001; ****, p<0.0001. Significance is shown only for cefsulodin, meropenem, and ampicillin ZOIs compared to Wild Type. All comparisons between ampicillin and ampicillin + sulbactam ZOI values are significant for each strain. A complete list of statistical comparisons between strains can be found in the supplemental table. D) Minimum inhibitory concentration (MIC) determined by MIC strips of indicated strains to ampicillin. Numbers on strips represent antibiotic concentration.

ExoA catalyzes the first committed step in succinoglycan biosynthesis from the precursor UDP-D-glucose and deletion of *exoA* blocks synthesis of succinoglycan and prevents cell spreading during treatment with β-lactam antibiotics (Glucksmann *et al*., 1993; Williams *et al*., 2022). The Δ*exoA* strain displayed an increase in sensitivity to all three antibiotics, as determined by a disc diffusion assay, but the degree of sensitivity does not fully explain the hypersensitivity of *chvG* or *chvI* deletion mutants (Figure 6B, Supplemental Table 10). Remarkably, a zone of succinoglycan accumulation is clearly visible in wildtype cells growing just beyond the zone of inhibition in the presence of cefsulodin and meropenem discs (Supplemental Figure 6B; blue arrows) suggesting that succinoglycan production is enhanced in these regions. In contrast, the accumulation of succinoglycan is not detectable in the presence of ampicillin discs, suggesting that the high degree of resistance to this drug may mask any contributions of succinoglycan to protection against this β-lactam. To remove β-lactamase-driven resistance mechanisms, we used disks containing both ampicillin and sulbactam, a β- lactamase inhibitor, and observed increased ampicillin sensitivity for Wild type, Δ*chvG*, Δ*chvI* and the Δ*exoA* mutants (Figure 6B, far right panel, Supplemental Table 10) confirming the contributions of β-lactamase to this resistance. Notably, the combined ampicillin and sulbactam treatment leads to the accumulation of succinoglycan in the region adjacent to the zone of inhibition in wild-type cells (Supplemental Figure 6B). Together, these results suggest that a ChvG-ChvI independent β-lactamase contributes to the resistance to ampicillin and that succinoglycan accumulation is beneficial if the β- lactamase is inhibited.

To better understand the contributions of both succinoglycan and β-lactamases to ampicillin resistance, we used MIC (minimum inhibitory concentration) strips to determine the sensitivity of our strains with more resolution (Figure 6C). Wild-type cells are highly resistant to ampicillin with a MIC greater than 96 μg/mL and deletion of either *chvG* or *chvI* greatly decreases the MIC to 4-6 μg/mL. Remarkably, loss of succinoglycan also greatly decreases the MIC to 6-8 μg/mL demonstrating that succinoglycan does contribute significantly to ampicillin resistance. When ampicillin and sulbactam are both present in the MIC strip, the MIC of wild-type cells to ampicillin drops to 16 μg/mL whereas the MIC for the Δ*exoA* mutant is only 2 μg/mL (Supplemental Figure 6C). These data suggest that a β-lactamase and accumulation of succinoglycan due to ChvG-ChvI activation are largely responsible for ampicillin resistance in *A. tumefaciens*.

The genome of *A. tumefaciens* encodes genes for ten β-lactamases (Figure 7A). Eight encode putative Ambler Class B metalo-β-lactamases (yellow), Atu0933 that encodes a putative Ambler Class D β-lactamase (purple), and *ampC* (Atu3077) that encodes a functional ortholog of the Ambler Class C β-lactamase AmpC (green). We were unable to find any genes encoding putative Ambler Class A β-lactamases (red) in the *A. tumefaciens* genome. Unsurprisingly, Δ*ampC* was highly sensitive to ampicillin as previously reported (Figueroa-Cuilan et al., 2022; Figure 7B; Supplemental Figure 7, Supplemental Table 10). There is a slight, but non-significant (p = 0.0627) increase in the sensitivity of the Δ*ampC* mutant to ampicillin in the presence of sulbactam, (Figure 7B, Supplemental Figure 7, Supplemental Table 10, Figueroa-Cuilan et al., 2022).

**Figure 7.**
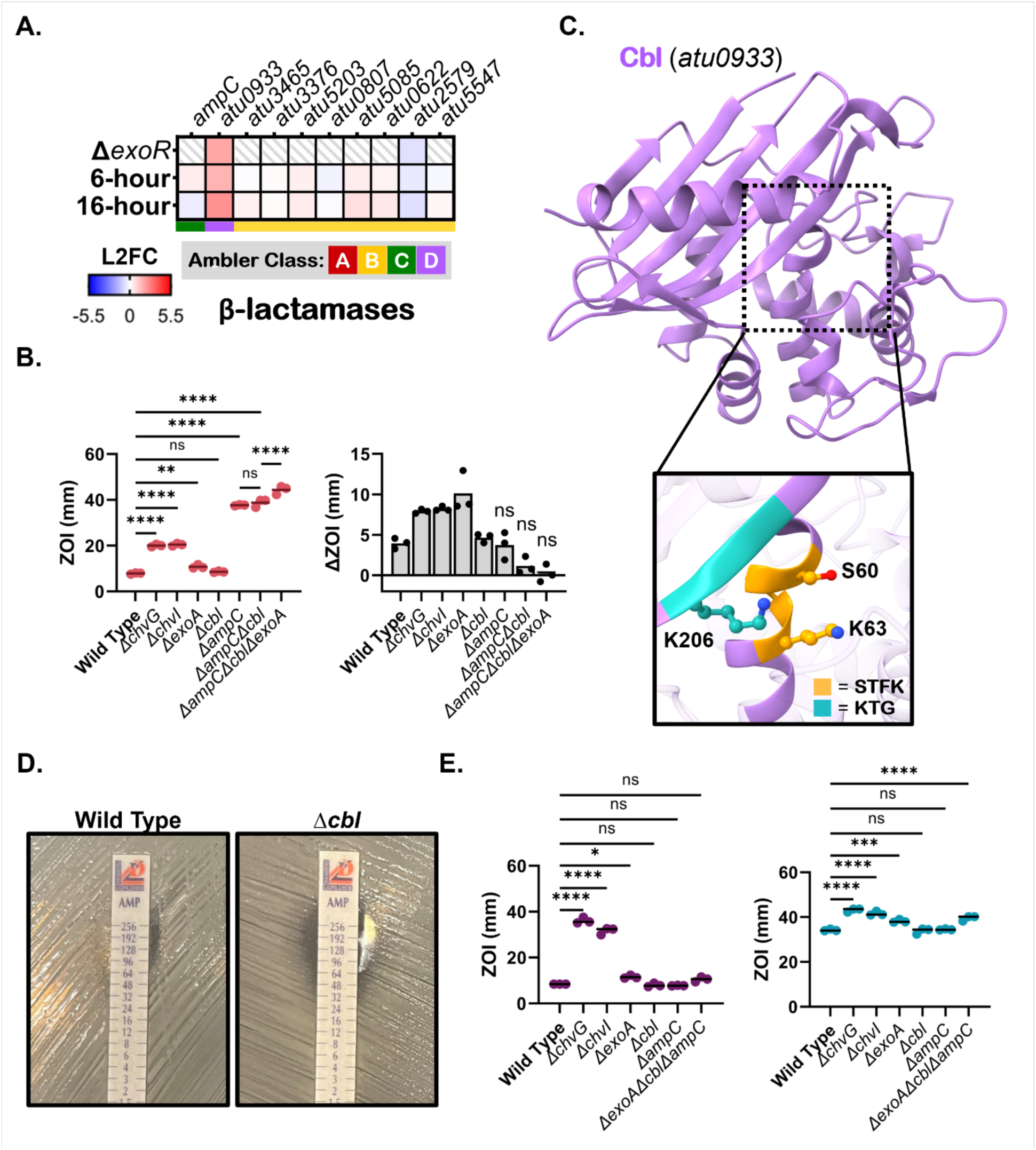
Contributions of ChvG-ChvI independent and dependant β-lactamases production on β-lactam resistance. A) Log 2-fold change (L2FC) in relative transcript abundance of ten predicted β-lactamase genes encoded by *Agrobacterium tumefaciens* in three ChvG-ChvI activating conditions, deletion of *exoR* (Heckel et al., 2014) and depletion of PBP1a for 6- or 16-hours (Williams et al., 2022). Shading represents L2FC where downregulated genes are a gradient of blue and upregulated genes are a gradient of red. White represents no change and white with gray stripes represents L2FC values that were insignificant and not reported. B) Disk diffusion assay with ampicillin (10 µg) measuring zone of inhibition (ZOI) for each strain. Change in ZOI from ampicillin to ampicillin + the β-lactam inhibitor sulbactam (10 µg ampicillin, 10 µg sulbactam) is show on the right. Comparison of means for ampicillin and ampicillin + sulbactam disks was performed with a two-way ANOVA, followed by Šídák’s multiple comparisons test with a single pooled variance. Comparisons for change in zone of inhibition were between ZOI of ampicillin + sulbactam and ZOI of ampicillin for the same strain. ns, not significant; *, p<0.1; **, p<0.01; ***, p<0.001; ****, p<0.0001. Only non- significant comparisons between ampicillin and ampicillin + sulbactam for each strain are shown. A complete list of statistical comparisons between strains can be found in the supplemental table. C) Crystal structure of Cbl (6NHU, 2.3 Å). Conserved Class D β-lactamase motifs shown in orange (STFK) and cyan (KTG). S60 is the putative active site. D) Minimum inhibitory concentration (MIC) determined by MIC strips of wildtype and **Δ***cbl* to ampicillin. Numbers on strips represent antibiotic concentration (µg/mL). E) Disk diffusion assays were conducted with cefsulodin (left, 100 µg) and meropenem (right, 10 µg) and the zones of inhibition (ZOI) for each strain are shown. Comparison of means for cefsulodin and meropenem disks was performed with a one-way ANOVA followed by Tukey’s multiple comparisons test with a single pooled variance. ns, not significant; *, p<0.1; **, p<0.01; ***, p<0.001; ****, p<0.0001.

These observations suggest the presence of another β-lactamase that may have minor contributions to ampicillin resistance.

We investigated ChvG-ChvI activated transcriptional datasets for a possible ChvG-ChvI regulated β-lactamase and found that transcription of Atu0933 is increased (Figure 7A, Yuan et al., 2008; Heckel et al., 2014; Williams et al., 2022), suggesting that expression of this putative β-lactamase may be induced during ChvG-ChvI activation. The amino acid sequence shows that the STFK and KTG motifs conserved in most Class D β- lactamase are present, and the crystal structure (6NHU) suggests that the residues of these motifs are oriented in a way that is indicative of a functional Class D β-lactamase (Figure 7C). Based on these pieces of evidence, we will henceforth refer to Atu0933 as ChvI-driven β-lactamase or Cbl. Structural alignments between Cbl and 61 other β- lactamase crystal structures using FoldTree clustered Cbl with other Class D β- lactamases such as the Cbl ortholog from *Agrobacterium tumefaciens* LBA4213 (6V6N), OXA-45 from *P. aeruginosa* (4GN2), OXA-427 from *Klebsiella pneumoniae* (6HUH), OXA-1 from *Escherichia coli* (3ISG), and CD0458 from *Clostridioides difficile 630* (6WY4) (Supplemental Figure 8A). Conservation of Cbl orthologs within the Hyphomicrobiales is somewhat erratic, an observation that is not altogether surprising given the tendency for genes encoding β-lactamases to be passed via horizontal gene transfer mechanisms often leading to strain-specific conservation. Presence of Cbl orthologs is primarily confined to the *Mesorhizobium* and *Agrobacterium* genera; however, we did find putative orthologs in two species of *Brucella* (*B. rhizosphaerae* and *B. gallinifaecis*) and one species of *Bartonella* (*B. tamiae)* (Supplemental Figure 8B).

Deletion of *cbl* did not significantly increase sensitivity to ampicillin in the disc diffusion assays, nor was there a notable additive effect in the *ΔampCΔcbl* double mutant (Figure 7B, Supplemental Table 10). However, this assay is not very sensitive, and we reasoned that it may not give us the resolution to observe weak effects. We therefore tested our strains using MIC strips and observed an increase in ampicillin sensitivity with the *Δcbl* mutant (Figure 7D). When ampicillin and sulbactam are present in the MIC strip the ampicillin sensitivity increases indicating that β-lactamase activity remains, likely due to the presence AmpC. Indeed, the *ΔampCΔcbl* double mutant is highly sensitive to ampicillin and the presence of sulbactam no longer leads to increased sensitivity to ampicillin (Supplemental Figure 7). This suggests that under these conditions AmpC is primarily responsible for β-lactamase activity, with Cbl potentially contributing a modest degree of secondary defense.

Next, we sought to determine if the contributions of AmpC and Cbl remain the same during challenges with cefsulodin and meropenem (Figure 7E, Supplemental Tables 11- 12). We find that *ΔampC* and *Δcbl* are resistant to both cefsulodin and meropenem suggesting that these responses are either specific for ampicillin or target other antibiotics. Finally, we constructed a triple mutant (Δ*exoA*Δ*ampC*Δ*cbl*) to assess the combined contributions of succinoglycan production and the β-lactamases AmpC and Cbl. The triple mutant displayed the greatest sensitivity to ampicillin out of all tested strains suggesting that ChvG-ChvI activation may make modest contributions to ampicillin resistance through succinoglycan and Cbl although the ampicillin resistance is primarily mediated by AmpC (Figure 7B; Figueroa-Cuilan et al., 2022). Remarkably, the triple mutant displayed similar resistance as the Δ*exoA* mutant when challenged with cefsulodin and meropenem (Figure 7E, Supplemental Table 11-12) indicating that succinoglycan confers protection against these drugs whereas neither the AmpC nor Cbl β-lactamases are protective. Overall, these results suggest that ChvG-ChvI activation is beneficial during challenge with β-lactam antibiotics in part because succinoglycan biosynthesis is upregulated.

### Activation of ChvG-ChvI may promote cell surface remodeling

Because succinoglycan production and Cbl expression do not fully explain the hypersensitivity of the *ΔchvG* and *ΔchvI* mutants, we continued to explore other ChvI regulon components that may help resist cell wall stress. Of the 14 LDTs encoded in the genome of *A. tumefaciens,* expression of four (*atu1164*, *atu3332*, *atu2133*, and *atu3331*) are transcriptionally upregulated under ChvG-ChvI activating conditions and *atu0844* is transcriptionally downregulated as compared to the wildtype transcript levels (Figure 8A). *A. tumefaciens* LDTs have been clustered into three distinct groups based on sequence similarity and structure (Aliashkevich *et al*., 2024). The absence of group 1 (*Δgr1*) or group 3 LDTs (*Δgr3*) results in reduced LD-crosslinking (Aliashkevich *et al*., 2024), thus we sought to determine if the absence of LDTs causes sufficient damage to activate the ChvG-ChvI pathway. We find that the LDT mutants retain the ability to produce compact microcolonies and that the ChvG-ChvI regulon can be activated in the LDT mutants by the presence of cefsulodin as indicated by cell spreading (Figure 8B). These observations are consistent with the fact that *ΔchvI* and *ΔchvG* mutants do not have significantly different muropeptide profiles compared to Wild type (Supplemental Figure 9A-B). Furthermore, phosphomimetic ChvI^D52E^ does not cause changes in the muropeptide profile and treatment with acid or cefsulodin only slightly increases the abundance of LD-crosslinks in the PG (Supplemental Figure 9A-B). Overall, these results suggest that the increased expression of LDTs during ChvG-ChvI activation likely does not primarily function to increase LD-crosslinking in PG.

**Figure 8.**
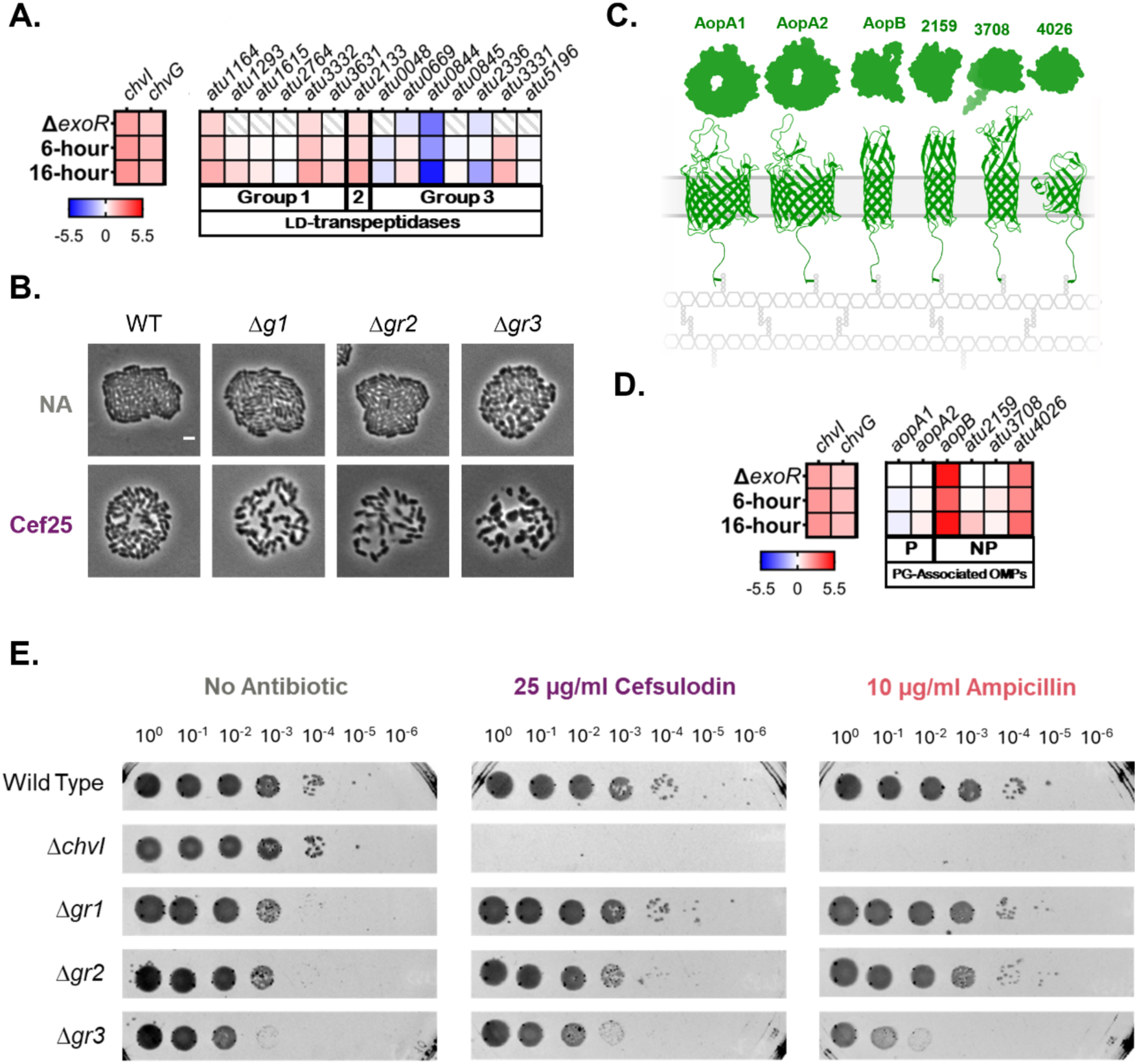
Differential expression of LD-transpeptidases and β-barrel outer membrane proteins during ChvG-ChvI activation may contribute to resistance to cefsulodin and ampicillin. A) Log_2_-fold change (L2FC) in relative transcript abundance of all 14 genes encoding LDTs under two ChvG-ChvI activating conditions, deletion of *exoR* and depletion of PBP1a for 6- or 16-hours. Shading represents L2FC where downregulated genes are a gradient of blue and upregulated genes are a gradient of red. White represents no change and white with gray stripes represents L2FC values that were insignificant and not reported. LDTs are grouped as described in Aliashkevich and Guest *et al*.(Aliashkevich et al., 2024). B) Microcolony production after 16 hours of growth for the Δ*gr1* (Δ*atu3631,* Δ*atu3332,* Δ*atu2764,* Δ*atu1615,* Δ*atu1293,* Δ*atu1164)*, Δ*gr2* (Δ*atu2133)* and Δ*gr3* (Δ*atu5196,* Δ*atu3331,* Δ*atu2336,* Δ*atu0845,* Δ*atu0844,* Δ*atu0669,* Δ*atu0048*) strains on a 1.5% ATGN agarose pad containing either no antibiotic or 25 µg/mL cefsulodin. Spreading microcolonies are indicative of succinoglycan production associated with ChvGI activation. Scale bar represents 2 µm. C) Alphafold-predicted structures of the 6 β-barrel OMPs in *A. tumefaciens* show that porins have a larger size. Both Top-down surface view and side ribbon view are shown for each protein.These β-barrel OMPs can link to the peptidoglycan (Sandoz et al., 2021). D) Log 2-fold change (Log2FC) in relative transcript abundance of genes encoding 6 β-barrel OMPs under ChvG-ChvI activating conditions as described for A. For β-barrel OMPs, predicted function as a porin (P) or not porin (NP) is indicated. E) Cell viability for each strain on ATGN plates containing no antibiotic, 25 µg/ml cefsulodin, or 10 μg/mL ampicillin is assessed by spotting serial dilutions incubated for 3 days.

The group 3 LDTs share similarity with the LDTs from *Brucella* which link outer membrane β-barrel proteins to PG (Supplemental Figure 9C, Godessart et al., 2020) and both group 1 and 3 LDTs are implicated in outer membrane β-barrel protein tethering to PG (Aliashkevich *et al*., 2024). Thus, we considered the possibility that β- barrel outer membrane proteins (OMPs) linked to PG by LDTs may provide structural integrity. Among the 6 potential LDT β-barrel substrates (Figure 8C; Sandoz et al., 2021), the transcripts for *aopB* and *atu4026* are strongly upregulated under conditions that activate ChvG-ChvI (Figure 8D). These β-barrels proteins are relatively small, in contrast to the much larger β-barrels, Atu1020 and Atu1021, which likely function as porins (Figure 8C). Based on these transcriptional findings, we hypothesize that LDT-mediated cross-linking of β-barrel proteins to the PG may confer a means to resist cell wall stress by remodeling the cell surface.

It is challenging to investigate the contributions of the individual LDTs due to functional redundancy (Aliashkevich *et al*., 2024). However, we reasoned that mutants lacking the LDT groups may inform us on their function within the ChvG-ChvI regulon. We then tested Δ*gr1*, Δ*gr2*, and Δ*gr3* mutants for growth viability on ATGN media containing no antibiotic, 25 µg/mL cefsulodin, or 10 µg/mL ampicillin. Δ*gr1* and Δ*gr2* mutants do not have increased sensitivity to either antibiotic; however, Δ*gr3* exhibits decreased viability to both cefsulodin and ampicillin (Figure 8E). Together these data indicate that ChvG-ChvI activation may change the outer membrane proteome through upregulation of transcripts encoding a subset of β-barrel proteins and LDTs. Outer membrane remodeling, combined with overproduction of succinoglycan, and activity of the Cbl β- lactamase confers protection under conditions that induce cell wall damage.

## Discussion

We previously found that inhibition of polar growth by depletion of PBP1a or treatment with the β-lactam antibiotic cefsulodin activates ChvG-ChvI in *A. tumefaciens* (Williams *et al*., 2022). We also demonstrated that a Δ*chvI* strain was hypersensitive to treatment with several β-lactam antibiotics including cefsulodin and ampicillin, suggesting a link between cell wall synthesis and the ChvG-ChvI two component system. To better understand the hypersensitivity of *ΔchvI* to cefsulodin, we used adaptive evolution to generate a suppressor strain of Δ*chvI* that displays restored wild-type levels of resistance to the antibiotic, but surprisingly not to ampicillin (Figure 1). Whole-genome sequencing revealed that the suppressor mutant had acquired two point mutations, one near the active site of PBP1a and another in the periplasmic tail of the outer-membrane protein AopA1. The PBP1a mutation partially restores growth to *ΔchvI* when challenged with cefsulodin but not ampicillin (Figure 2). Given that treatment with cefsulodin and ampicillin produce distinct morphological phenotypes, we hypothesize that cefsulodin targets PBP1a and polar elongation whereas ampicillin blocks cell division. We find that a phosphomimetic ChvI is sufficient to restore growth in the presence of both cefsulodin and ampicillin and that the *chvG*-*chvI* promoter is activated by the presence of an array β-lactam and other cell wall targeting antibiotics (Figure 3). These observations led us to speculate that ChvG-ChvI is likely to be activated by accumulation of cell wall damage, rather than loss of a specific enzymatic activity.

During the depletion of PBP1a, secretion of the exopolysaccharide succinoglycan is readily detectable due to the flagellar-independent cell surface spreading phenotype (Williams *et al*., 2022). In contrast, many *A. tumefaciens* cell division mutants fail to divide and form microcolonies, which makes the cell spreading phenotype more difficult to assess (Howell *et al*., 2019; Williams *et al*., 2021). Here we confirm that ChvG-ChvI activation occurs in response to depletion of essential cell division proteins causing overproduction of succinoglycan (Figure 4). Remarkably, depletion of FzlA, an FtsZ-binding protein that coordinates activation of sPG synthesis and chromosome segregation, causes cell spreading (Figure 5; Lariviere et al., 2019; Payne et al., 2024). In support of our hypothesis that disrupted septal PG synthesis within the SEDS-bPBP complex is leading to ChvG-ChvI activation, a single hyperactivating mutation in *A. tumefaciens* FtsW (F137L) is sufficient to restore microcolony formation, prevent cell spreading, and restore basal expression of the *chvG-chvI* promoter during *fzlA* depletion (Figure 5; Lariviere et al., 2019). Together, these results illustrate the non-specific nature of ChvG-ChvI activation and suggest that ChvG-ChvI responds broadly to conditions that result in impaired cell wall synthesis or the accumulation of cell wall damage.

Orthologs of ChvG and ChvI have been described in *Brucella abortus* (BvrS-BvrR), *Bartonella henselae* (BatS-BatR), *Rhizobium leguminosarum* (ChvG-ChvI), and *Caulobacter crescentus* (ChvG-ChvI)(Charles and Nester, 1993; Cheng and Walker, 1998; Lamontagne *et al*., 2007; Quebatte *et al*., 2010; Vallet *et al*., 2020). Reports of activation of ChvG-ChvI orthologs by cell wall synthesis inhibition, disruptions to membranes, and osmotic stress have all been proposed to activate ChvG-ChvI. Together, these reports suggest a common, conserved function for ChvG-ChvI and its orthologs during cell envelope stress within the Alphaproteobacteria.

Though these commonalities among orthologs are expanding our understanding of the conditions that activate the ChvG-ChvI pathway, we still have a limited understanding of how the ChvG-ChvI regulon is protective against cell surface stressors. Here, we explored the contributions of ChvG-ChvI-mediated succinoglycan production, β- lactamase activity, and cell surface remodeling to survival during cell wall stress.

First, we found that the biosynthesis of the exopolysaccharide succinoglycan plays a role in ChvG-ChvI-conferred resistance to ampicillin, cefsulodin, and high concentrations of meropenem (Figure 6). Succinoglycan is an acidic exopolysaccharide synthesized and secreted by plant-associated Rhizobia (Cangelosi *et al*., 1987; Glucksmann *et al*., 1993; Wu *et al*., 2016). The function of succinoglycan in *A. tumefaciens* remains obscure, but it is required for symbiosis in *S. meliloti* and has been shown to be protective against hydrogen peroxide-dependent damage and the plant-derived antimicrobial peptide NCR247 (Lehman and Long, 2013; Arnold *et al*., 2017, 2018). While the chemical mechanisms of protection are unknown, overproduction of succinoglycan is not sufficient to provide complete protection against NCR247 leading to the hypothesis that it may form a proximal protective layer around the bacteria (Arnold *et al*., 2018). Our observation that succinoglycan contributes to tolerance to multiple β-lactam antibiotics is consistent with the possibility that this exopolysaccharide forms a barrier around cells to promote survival in an array of stressful conditions.

Next, we found that expression of Cbl, an Ambler Class D β-lactamase was upregulated during ChvG-ChvI activation (Figure 7). Notably, we found that Cbl only had a modest contribution to ampicillin resistance and does not contribute to resistance to cefsulodin or meropenem. However, Cbl may play an auxiliary role in ampicillin resistance which is masked by the primary defense, an ampicillin-targeting Ambler Class C β-lactamase, AmpC (Figueroa-Cuilan *et al*., 2022) In addition, it is likely that Cbl has additional β- lactam targets not tested here. Cbl is structurally very similar to several oxacillinase (OXA) enzymes and therefore may function in hydrolyzing oxacillin (Supplemental Figure 8). Cbl is structurally similar to OXA-45 from *Pseudomonas aeruginosa* which is an extended spectrum-OXA that can hydrolyze cephalosporins such as ceftazidime, cefotaxime, and aztreonam but does not target carbapenems or cephamycins (Toleman *et al*., 2003). It will be of interest to further characterize the substrate profile of Cbl to determine if it contributes to resistance to other β-lactam antibiotics. Proprietary blends of β-lactam and β-lactamase inhibitors such as Timentin and Augmentin are routinely used to clear *Agrobacterium* following plant transformation (Cheng et al., 1998; Ieamkhang and Chatchawankanphanich, 2005; Morton and Fuqua, 2012). Thus, an improved understanding of mechanisms of contributing to the high level of β-lactam resistance should enable engineering of strains which remain competent for transformation but are easier to clear following plant transformation.

Finally, since LD-transpeptidase activity is known to promote β-lactam resistance (Mainardi *et al*., 2000; Hugonnet *et al*., 2016) and the *aopB* transcript, which encodes a β-barrel outer membrane protein, is significantly upregulated when ChvG-ChvI is activated (Figure 8; Heckel et al., 2014; Williams et al., 2022) we considered the contributions of cell surface remodeling to survival under cell wall stress. Although elongation inhibition via cefsulodin treatment (Supplemental Figure 9A-B) or PBP1a depletion (Heckel *et al*., 2014; Williams *et al*., 2022) increases the proportion of LD-crosslinks in peptidoglycan and a subset of transcripts encoding LDTs are impacted by ChvG-ChvI activation, there is no evidence to suggest that the ChvG-ChvI activation causes significant changes to the peptidoglycan composition (Supplemental Figure 9A- B). Notably, expression of the genes encoding AopB and its orthologs (known as RopB) are upregulated during ChvG-ChvI activation in *A. tumefaciens* (Li *et al*., 2002; Heckel *et al*., 2014; Alakavuklar *et al*., 2021; Williams *et al*., 2022), *S. meliloti* (Chen *et al*., 2009), *R. leguminosarum* (Foreman *et al*., 2010), *B. henselae* (Quebatte *et al*., 2010), and *Brucella abortus* (Guzmán-Verri *et al*., 2002). AopB is attached to peptidoglycan via crosslinking mediated by LDTs (Godessart *et al*., 2020; Sandoz *et al*., 2021) raising the possibility that linking AopB to the peptidoglycan may be increased to confer structural stability during cell wall stress. Although *ΔaopB* does not have an obvious growth defect or change in peptidoglycan composition, AopB tethering to peptidoglycan is increased in the *Δgr3* LDT mutant when grown in LB with 5% salt which induces morphological changes (Aliashkevich *et al*., 2024) consistent with the possibility that increased AopB tethering may provide structural integrity to prevent lysis. These data are consistent with observations in *Rhizobium leguminosarum* where RopB functions to support outer membrane integrity as *ropB* mutants are more sensitive to detergents, hydrophobic antibiotics, and a weak organic acid (Foreman *et al*., 2010). Furthermore, hypersensitivity of *ΔchvG* to these stressors was bypassed by constitutive *ropB* expression (Vanderlinde and Yost, 2012). Alternatively, RopB adopts an amyloid fibril structure which is observed in the capsular polysaccharides of stationary phase free- living cells and promotes host-symbiont interactions (Kosolapova *et al*., 2019, 2022).

Bacterial amyloid proteins often function as a structural scaffold in biofilms and enhance antibiotic resistance (Akbey and Andreasen, 2022). Thus, it is intriguing to speculate that the *A. tumefaciens* AopB may be capable of adopting an amyloid state and, along with succinoglycan, contribute to the formation of a protective barrier against cell wall targeting antibiotics. Notably, the second mutation identified in our SNP analysis of the cefsulodin tolerant *ΔchvI* is in located in the AopA1 periplasmic tail near the site essential for crosslinking to peptidoglycan (Figure 1B, Supplemental Figure 1) providing additional support for a role in outer membrane remodeling and/or amyloid production during cell wall stress in *A. tumefaciens*. Elucidating the outer membrane remodeling that occurs in *A. tumefaciens* cells with cell wall damage is an important focus of future research that will provide key insights into the function of ChvG-ChvI activation and β- lactam antibiotic mechanisms in this important phytopathogen.

## Materials and Methods

### Bacterial strains, plasmids, and growth conditions

A list of all bacterial strains and plasmids used in this study is provided in the Supplemental Table 15. *Agrobacterium tumefaciens* C58 and derived strains were grown in ATGN minimal media (Morton and Fuqua, 2012b) without exogenous iron as stated at 28°C with shaking. When appropriate, kanamycin (KAN) was used at the working concentration of 300 μg/ml.

When indicated, isopropyl β-D-1-thio-galactopyranoside (IPTG) was used as an inducer for depletion strains at a concentration of 1 mM. *E. coli* DH5α and S17-1 λ pir were grown in Lysogeny Broth medium at 37°C and when appropriate 50 μg/ml or 30 μg/ml of KAN were added, respectively.

### Construction of plasmids and strains

Vectors for allelic exchange to generate PBP1a^V659M^ were constructed using recommended methods for *A. tumefaciens (Morton and Fuqua, 2012a)*. Primer sequences and additional cloning details are described in Supplemental Table S1. Briefly, the *mrcA* sequence was amplified using P1 and P4 and digested and ligated into *sacB* suicide plasmid pNPTS139, then confirmed by sequencing. From this vector source, primer pairs M13F and P2, and P3 with P4 were used in separate reactions to amplify the first and second halves of *mrcA* up to the codon that encodes amino acid residue 659, with P2 and P3 each including the desired mutation in the center of their sequence. Amplicons were spliced together by a SOEing reaction using primer pair M13F/P4. The amplicon was digested with BamHI and SpeI and ligated with T4 ligase into pNTPS139. This deletion plasmid was introduced into *A. tumefaciens* by mating using an *E. coli* S17 conjugation strain to generate kanamycin-resistant and sucrose-sensitive primary integrates. Primary integrates were grown overnight in media with no selection. Secondary recombinants were screened by patching for sucrose resistance and kanamycin sensitivity. These were further screened for the point mutation phenotype by 16-hour overnight microscopy on a 1.5% ATGN agarose pad containing 25µg/ml cefsulodin. Microcolonies that failed to spread in WT or survived in Δ*chvG*/Δ*chvI* strains were considered phenotypic hits. Colony PCR with primers P5/P6 for the respective sequence target was used to amplify and subsequently sequenced to further confirm allelic replacement of the gene.

pMR15 P_chvGI_-*lacZ* was constructed by amplifying the 338 bp intergenic region just upstream of the *chvI-chvG-hprK* operon of *Agrobacterium tumefaciens* with primers flanked with the restriction digestion sites PstI and NheI. Restriction digestion of the pMR15-*lacZ* and the intergenic region amplicon was performed, and the fragments were ligated and transformed as described above. Vector was introduced to each *A. tumefaciens* strain using the S17 *E. coli* as a conjugal donor and vector presence was confirmed by PCR.

### Cefsulodin adaptive evolution experiment

Wildtype and **Δ***chvI* colonies were inoculated in 1 mL of ATGN media and grown overnight at 28°C in a shaking incubator. Overnight culture was diluted to an OD_600_ of 0.1 and grown to an OD_600_ of ∼0.7. The cultures were diluted to an OD_600_ of 0.05 split into two 500 µL cultures in a 48 well plate. To select **Δ***chvI* suppressors, cefsulodin was added to one well of the **Δ***chvI* culture to a final concentration of 4 µg/mL. Cultures were grown overnight and each morning were diluted 1:10 in fresh media while the remainder was saved in a frozen stock. WT and control cultures of Δ*chvI* were also passaged without antibiotic. Optical densities (600 nm) of the overnight and freshly diluted cultures were obtained by a BioTek Synergy H1 Hybrid plate reader to monitor growth. Cefsulodin concentrations were increased by 25% each subsequent day where turbid growth occurred for 14 passages, reaching a maximum concentration of 75 µg/ml. Turbid cultures from the 14^th^ passage were saved at -80°C then stuck purified on ATGN media containing 50 µg/ml cefsulodin. Three isolates were cultured in media containing 25 µg/ml cefsulodin and the DNA of two isolates was successfully extracted using a Qiagen Blood & Cell Culture DNA mini Kit (13323) and submitted for whole genome sequencing at SeqCenter (https://www.seqcenter.com/). In addition, the DNA from three isolated colonies from wildtype and **Δ***chvI* prior to passaging (P0) and after 14 passages (P14) were extracted as described and submitted for whole genome sequencing to serve as controls for accumulation of non-specific mutations. Sequencing was performed as 150 bp paired end libraries on an Illumina ######.

Whole genome single nucleotide polymorphism (SNP) calling was performed as previously described (Weisberg *et al*., 2020). Briefly, BWA v. 0. mem with the parameter “-M” was used to map reads to the *Agrobacterium fabrum* C58 reference genome (NCBI: GCF_000092025.1)(Li and Durbin, 2009). Picard tools v.2.20 was used to mark duplicate reads and sort the alignments (Picard Tools - By Broad Institute). GATK v. 3.7 HaplotypeCaller with the parameters “-ERC GVCF -ploidy 1” was used to call partial variants for each sample . The GATK tool GenotypeGVCFs with the default parameters was used to call variants across all samples. The GATK tool SelectVariants was used to subset SNP variants, and the tool VariantFiltration with the parameters “-- filterExpression ’QD < 2.0’ --filterName ’QD2’ --filterExpression ’SOR > 3.0’ --filterName ’SOR3’ --filterExpression ’QUAL < 30.0’ --filterName ’QUAL30’ --filterExpression ’FS > 60.0’ --filterName ’FS60’ --filterExpression ’MQ < 40.0’ --filterName ’MQ40’ -- filterExpression ’MQRankSum < -12.5’ --filterName ’MQRankSum-4’ --filterExpression ’ReadPosRankSum < -8.0’ --filterName ’ReadPosRankSum-8’” was used to filter SNPs for quality. SnpEff v.4.3t with the C58 reference genome and parameters “-no-downstream -no-upstream -no-intron” was used to predict functional effects of each SNP (Cingolani *et al*., 2012).

### Growth curves

Exponential phase cells (OD_600_ 0.2 – 0.5) were diluted to an OD of 0.05 in ATGN media in a 96 well microplate to incubate at 28°C, orbitally shaking in a BioTek Synergy H1 Hybrid plate reader with OD_600_ readings taken every 10 minutes for up to 72 hours where indicated. Conditions included ATGN supplemented with 25 µg/ml cefsulodin or meropenem at concentrations of 0.75, 1.5, and 3.0 µg/ml. Growth curves were graphed using GraphPad Prism 10.3.1.

### AlphaFold2 predictive structural modeling and docking predictions

Structural predictions were done using AlphaFold2 (Yang *et al*., 2023). Visualization and electrostatic surface calculations of PBP1a and PBP1a* were done using UCSF ChimeraX 1.8. Docking of cefsulodin onto the AlphaFold2 predicted structure of PBP1a and PBP1a* was performed using AutoDock VINA (Trott and Olson, 2010) through the PyRx software (Dallakyan and Olson, 2015; Kondapuram *et al*., 2021).

### Phase and fluorescence microscopy

A small volume (∼0.4-1 μl) of exponential phase cells (diluted to an OD_600_ of 0.05) are applied to 1.5% low-melting point agarose pads made with either ATGN alone, with 200 mM calcofluor, with 25 µg/ml cefsulodin, or 10 µg/ml ampicillin respectively. When appropriate agarose pads were also supplemented with 1mM IPTG. Sealed coverslips with air channels were incubated overnight at 28°C for 16-24 hours before imaging.

Phase contrast and epifluorescence microscopy were performed with either an inverted Nikon Eclipse TiE or a Leica DMi8 inverted microscope equipped with a 63× 1.4 NA Plan Apochromat oil-immersion phase-contrast objective and a high precision motorized stage (Pecon). Excitation light was generated by a Lumencor Spectra-X multi-LED light source with integrated excitation filters. An XLED-QP quadruple-band dichroic beam- splitter (Leica) was used (transmission: 415, 470, 570, and 660 nm) with an external filter wheel for all fluorescent channels. Calcofluor-stained succinoglycan was imaged using the DAPI filter with an exposure time of 40 ms.

### Spotting assays

Exponential phase cells were diluted to an OD_600_ of 0.015 (OD_600_ obtained on a BioTek Synergy 96 well plate) in ATGN, then serially diluted ten-fold out to a 10^-6^ dilution factor in ATGN media before 4 µl were spotted onto agar plates. Where indicated, plates contained ATGN alone, 25 µg/ml cefsulodin, or 10 µg/ml ampicillin. Ampicillin plates were used for this assay within 2-5 days of generation. Once spots dried, plates were incubated for 3 days before imaging except in certain cases where incubation was carried out to 5 days (*chvI*^D52E^, Figure 3A).

### Miller assays

Ten 1 mL cultures of wild-type cells carrying the pMR15 P_chvGI_-*lacZ* plasmid were grown up overnight in 300 ug/mL of kanamycin for selection, back-diluted and then grown to exponential phase at an approximate OD_600_ of 0.5. Cultures were normalized to the lowest common concentration (approximately 0.4-0.5) then supplemented with corresponding antibiotic or water to achieve a concentration of 20 µg/ml except where indicated. Cells were grown in the presence of the antibiotic for 6 hours at 28°C. Depletion strains were grown overnight with 1mM IPTG and 300 µg/ml kanamycin, then washed of IPTG three times and cultures diluted to an OD of 0.1-0.2 then incubated at 28°C for 6 hours. For depletion strains with extended incubation, 100 µl of the IPTG-depleted cells was transferred to a new tube containing 1.9 ml ATGN media and incubated shaking overnight at 28°C. This was repeated the next day for strains incubating for up to 48hrs. After each respective incubation period, cultures were centrifuged at 4000 rpm for 10 minutes, supernatant discarded, then resuspended in water. OD600 of cultures was obtained and samples were diluted down either to match the OD_600_ of the least concentrated culture or to an OD of 0.6. The Miller assays were performed in a BioTek Synergy H1 Hybrid plate reader using a protocol described previously (Schaefer *et al*., 2016). BugBuster (Millipore; 70921-4) was substituted for PopCulture reagent.

### Disc diffusion assays

Exponential phase cells were diluted to an OD_600_ of 0.125 (OD_600_ obtained on a BioTek Synergy plate reader in a 96 well plate). A sterile cotton swab was soaked in the normalized culture then lawned onto an ATGN plate. Discs were placed sterilely onto the dried lawn with ∼0.75-1” of space between each disc and no more than two discs per lawn. Manufactured antibiotic discs were obtained from Thermo Scientific Oxoid while cefsulodin discs were generated by soaking sterile blank discs with 20 µl of cefsulodin stock to obtain a total of 100 µg per disc. Discs were dried for 2-3 hours before placement onto lawns. Plates were incubated for two days at 28°C and then imaged. Zones of inhibition were measured using NIH ImageJ software. Three independent replicates were conducted for each strain. For meropenem and cefsulodin discs, an ordinary one-way analysis of variance (ANOVA) was used to compare means in zone of inhibition diameter between strains. For ampicillin and ampicillin + sulbactam discs, a two-way analysis of variance (ANOVA) was used to compare the means between strains and between ampicillin and ampicillin + sulbactam discs.

### Ampicillin MIC strip assays

Exponential phase cells were diluted to an OD_600_ of 0.125 (OD_600_ obtained on a BioTek Synergy plate reader in a 96 well plate). A sterile cotton swab was soaked in the normalized culture then lawned onto an ATGN plate. MIC strips from Liofilchem placed in the center of the plate. Plates were incubated for two days at 28°C and then imaged. MIC concentrations were determined as the lowest concentration indicated on the strip inside of the ZOI.

### Extraction of sacculi and PG analysis

Extraction of sacculi were performed as previously described (Cingolani *et al*., 2012). Briefly, cells were grown overnight in 2.5 ml ATGN then added to 250 ml of media, a 1:100 dilution, and incubated at 28°C until they reached an OD_600_ between 0.5 – 0.8. Conditions included cefsulodin supplemented at a concentration of 25 µg/ml and ATGN at a pH of 5.5, buffered by 2-(N-morpholino)ethanesulfonic acid (MES). The cells were then centrifuged at 5250 x g for 10 mins at room temperature, supernatant discarded and resuspended in 3 ml of 1X PBS. The resuspensions were added to 6 ml of boiling 6% SDS, boiled for 3 hours with simultaneous mixing, then mixing continued overnight without heat.

The insoluble fraction (peptidoglycan) was pelleted at 400,000 x g, 15 min, 30 °C (TLA- 100.3 rotor; OptimaTM Max ultracentrifuge, Beckman) and resuspended in Milli-Q water.

This step was repeated 4-5 times until the SDS was washed out. Next, peptidoglycan was treated with Proteinase K 0.1 mg/ml at 37 °C for 1 h and then boiled in 1% SDS for 2 h to stop the reaction. After SDS was removed as described previously, peptidoglycan samples were resuspended in 200 μL of 50 mM sodium phosphate buffer pH 4.9 and digested overnight with 30 μg/ml muramidase (from Streptomyces albus) at 37 °C. Muramidase digestion was stopped by heat-inactivation (boiling during 5 min). Coagulated protein was removed by centrifugation (20,000 x g, 15 min). The supernatants (soluble muropeptides) were subjected to sample reduction. First, pH was adjusted to 8.5-9 by addition of borate buffer 0.5 M pH 9 and then muramic acid residues were reduced by sodium borohydride treatment (NaBH4 10 mg/ml final concentration) during 30 min at room temperature. Finally, pH was adjusted to 2.0-4.0 with orthophosphoric acid 25% prior to analysis by LC.

Chromatographic analyses of muropeptides were performed by Ultra Performance Liquid Chromatography (UPLC) on an UPLC system (Waters) equipped with a trapping cartridge precolumn (SecurityGuard ULTRA Cartridge UHPLC C18 2.1 mm, Phenomenex) and an analytical column BEH C18 column (130 Å, 1.7 μm, 2.1 mm, Waters) maintained at 45 °C. Muropeptides were detected by measuring the absorbance at 204 nm using an ACQUITY UPLC UV−visible Detector. Muropeptides were separated using a linear gradient from buffer A (Water + 0.1 % (v/v) formic acid) to buffer B (Acetonitrile 100 % (v/v) + 0.1 % (v/v) formic acid) over 15 min with a flowrate of 0.25 ml/min. The QTOF instrument was operated in positive ion mode, with data collection performed in untargeted MS^e^ mode. The parameters were set as follows: capillary voltage 3.0 kV, source temperature 120 °C, desolvation temperature 350 °C, sample cone voltage 40 V, cone gas flow 100 L h^−1^ and desolvation gas flow 500 L h^−1^. Mass spectra were acquired at a speed of 0.25 s/scan. The scan was in a range of 100–2000 m/z. Data acquisition and processing was performed using MassLynx or UNIFI software package (Waters Corp.). The quantification of muropeptides was based on their relative abundances (relative area of the corresponding peak) and relative molar abundances. A table of all the identified muropeptides and the observed ions is provided (Supplemental Table 13).

### Generation of Heatmaps

Log_2_ fold change heatmaps of transcriptional changes of succinoglycan, β-lactamase, and LD-transpeptidase genes were generated using GraphPad Prism 10.3.1. Similarity matrix of the 14 *Agrobacterium tumefaciens* LDTs and 8 *Brucella abortus* LDTs based on sequence similarity using Geneious Prime 2024.0.5 (https://www.geneious.com). Sequence similaritires were clustered and visualized as a heatmap using the Clustergrammer web app (Fernandez *et al*., 2017).

## Supporting information

Supplemental Figures 1-9 and Supplemental Reference

## Acknowledgements.

We acknowledge Amara Mason for technical assistance early during this work. We thank Amelia Randich, Brody Aubry, and all members of the Brown Lab for critical feedback on this manuscript.

