## Supplemental Figures 1-9 and Supplemental Reference for "Activation of the ChvG-ChvI pathway promotes multiple survival strategies during cell wall stress in *Agrobacterium tumefaciens*"

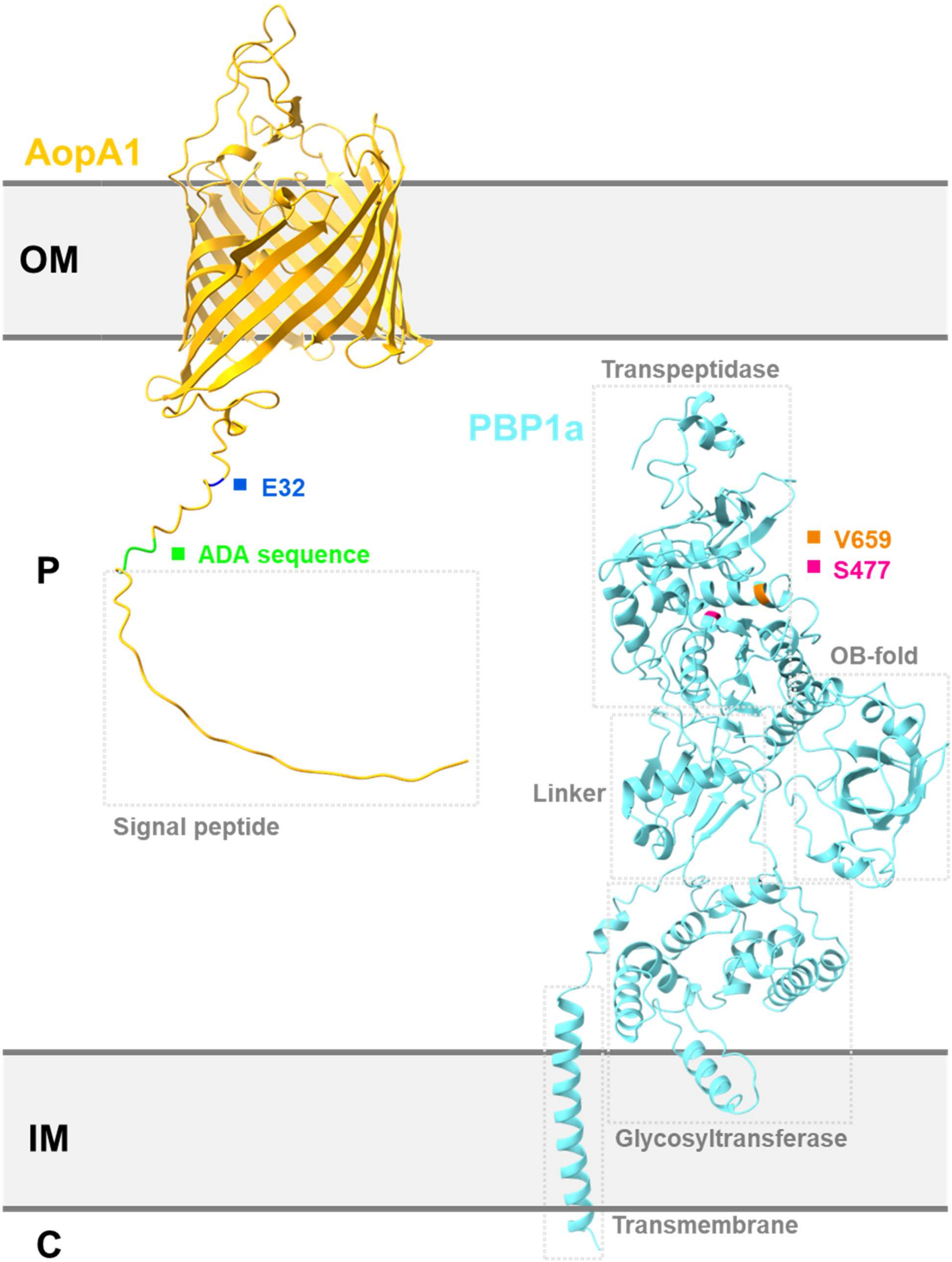
AopA1
OM
Transpeptidase
PBP1a
E32
V659
S477
ADA sequence
P
OB-fold
Linker
Signal peptide
Glycosyltransferase
IM
Transmembrane
C

**Supplemental Figure 1. Alphafold-predicted three-dimensional structures of AopA1 and PBP1a with respect to cellular topology.** The wildtype residues with mutations in Δ*chvI*$^{cef}$ are labeled for each (Blue E32 in AopA1 and orange V659 in PBP1a). The ADA sequence used for ld-transpeptidase mediated crosslinking of in AopA1 to the PG and PBP1a catalytic residue (S477) are highlighted in green and pink respectively. Structural and functional domains outlined in gray dotted boxes.

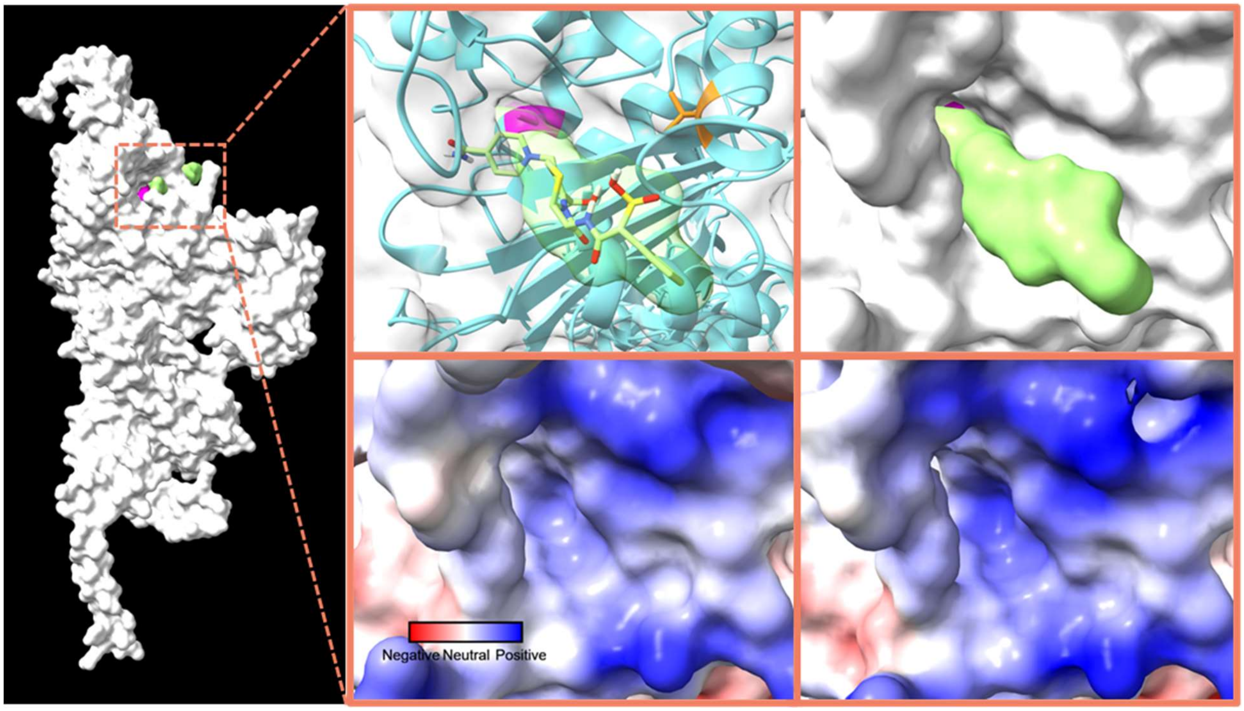


**Supplemental Figure 2. Alphafold predicts cefsulodin interaction with the wildtype PBP1a transpeptidase catalytic site and increased positive charge at the site in the Δ*chvI*$^{cef}$ mutant.** The cefsulodin molecule in green is predicted to interact with the transpeptidase catalytic site of PBP1a in WT as shown in a linear (A) and space-filling model (B). The catalytic residue for transpeptidation is labeled in panel A as pink and the valine residue that is mutated in Δ*chvI*$^{cef}$ is labeled in orange. C) Displays the electrical model of the transpeptidase catalytic site in WT. Red indicates a more negative charge, white is neutral, and blue indicates a more positive charge. D) Displays the same catalytic site as predicted in the Δ*chvI*$^{cef}$ strain with increased positive charge as a result of the methionine residue replacing valine.

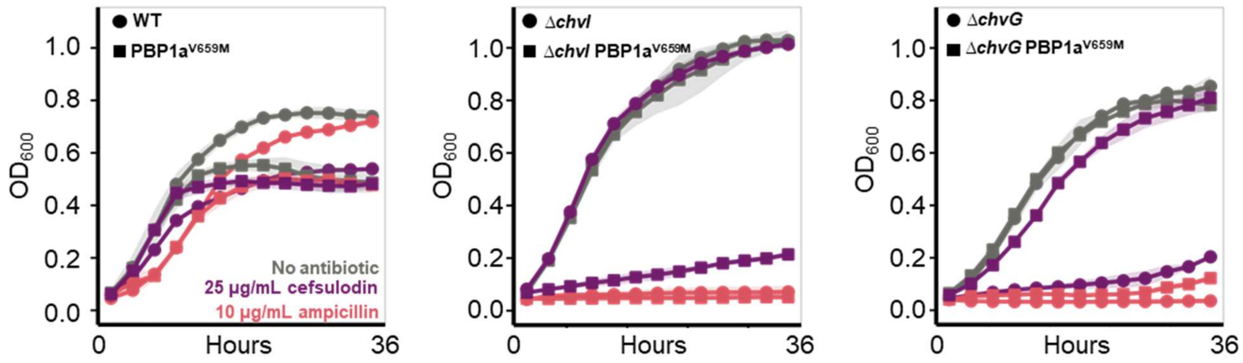


**Supplemental Figure 3. Cefsulodin-suppressing substitution rescues growth during treatment with 25 µg/mL cefsulodin in both Δ*chvI* and Δ*chvG* backgrounds.** Growth ($OD_{600}$) of Wild Type (WT), Δ*chvI,* and Δ*chvG* with (circles) and without (squares) the cefsulodin-suppressing PBP1a$^{V659M}$ substitution. Growth is shown in ATGN (no antibiotic, gray), ATGN supplemented with 25 µg/mL cefsulodin (purple), or ATGN supplemented with 10 µg/mL ampicillin (pink) measured over 36 hours. Data shown represents the average of 3 biological replicates, with shaded regions indicating standard deviation.

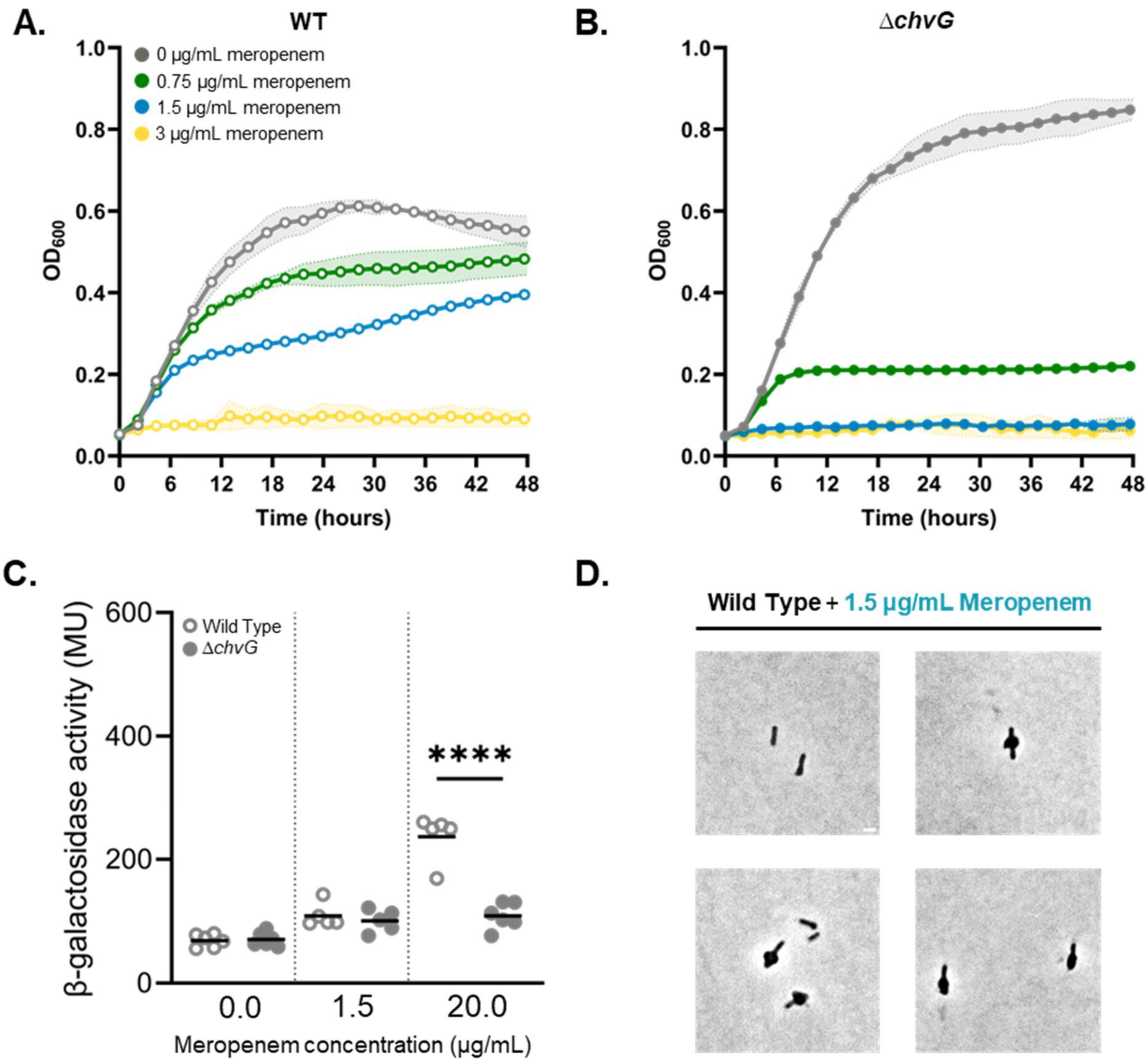


**Supplemental Figure 4. Treatment with minimum inhibitory concentration of meropenem does not activate ChvG-ChvI.** A and B) Growth curves for WT *Agrobacterium tumefaciens* and Δ*chvG* in ATGN media for 48 hours in no treatment, 0.75 µg/ml, 1.5 µg/ml, and 3 µg/ml meropenem. C) β-galactosidase activity in Miller Units (MU) under *chvGI* promoter after antibiotic treatment of exponential-phase WT and Δ*chvG* strains. Cells were incubated for 6 hours with meropenem at concentrations of 1.5 µg/ml (MIC) and 20 µg/ml. Significance compares each treatment to the no treatment control and was calculated with an ANOVA and Tukey's post hoc test. ****, $p < 0.00005$. Nonsignificant comparison bars were not shown. D) Phase micrographs of wild-type cells grown overnight on 1.5% ATGN agarose pads containing 1.5 µg/mL meropenem.

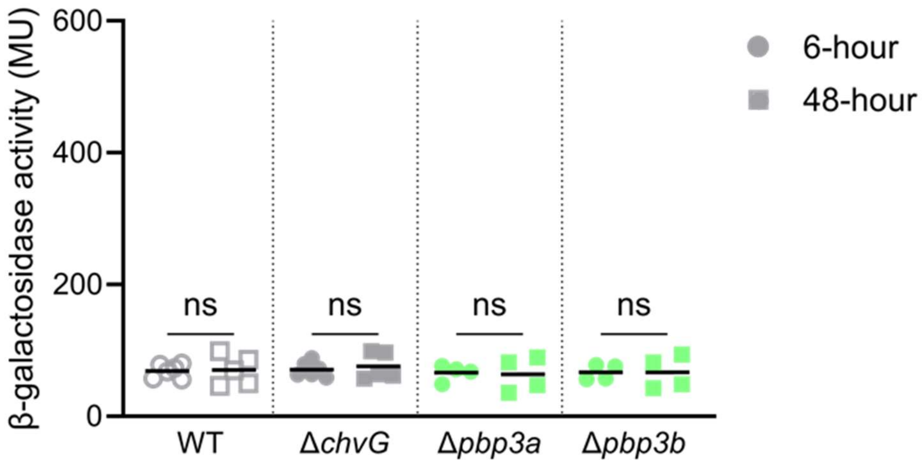


**Supplemental Figure 5. Single deletions of *pbp3a* and *pbp3b* do not activate ChvG-ChvI.** β-galactosidase activity in Miller Units (MU) as a reporter of *chvGI* promoter activity in wildtype Δ*chvG,* Δ*pbpb3a,* and Δ*pbp3b* strains at 6 and 48 hours of growth. Significance was calculated using t-tests for comparison of means between 6-hour and 48-hour time points of each strain. ns indicates that the difference are not significant.

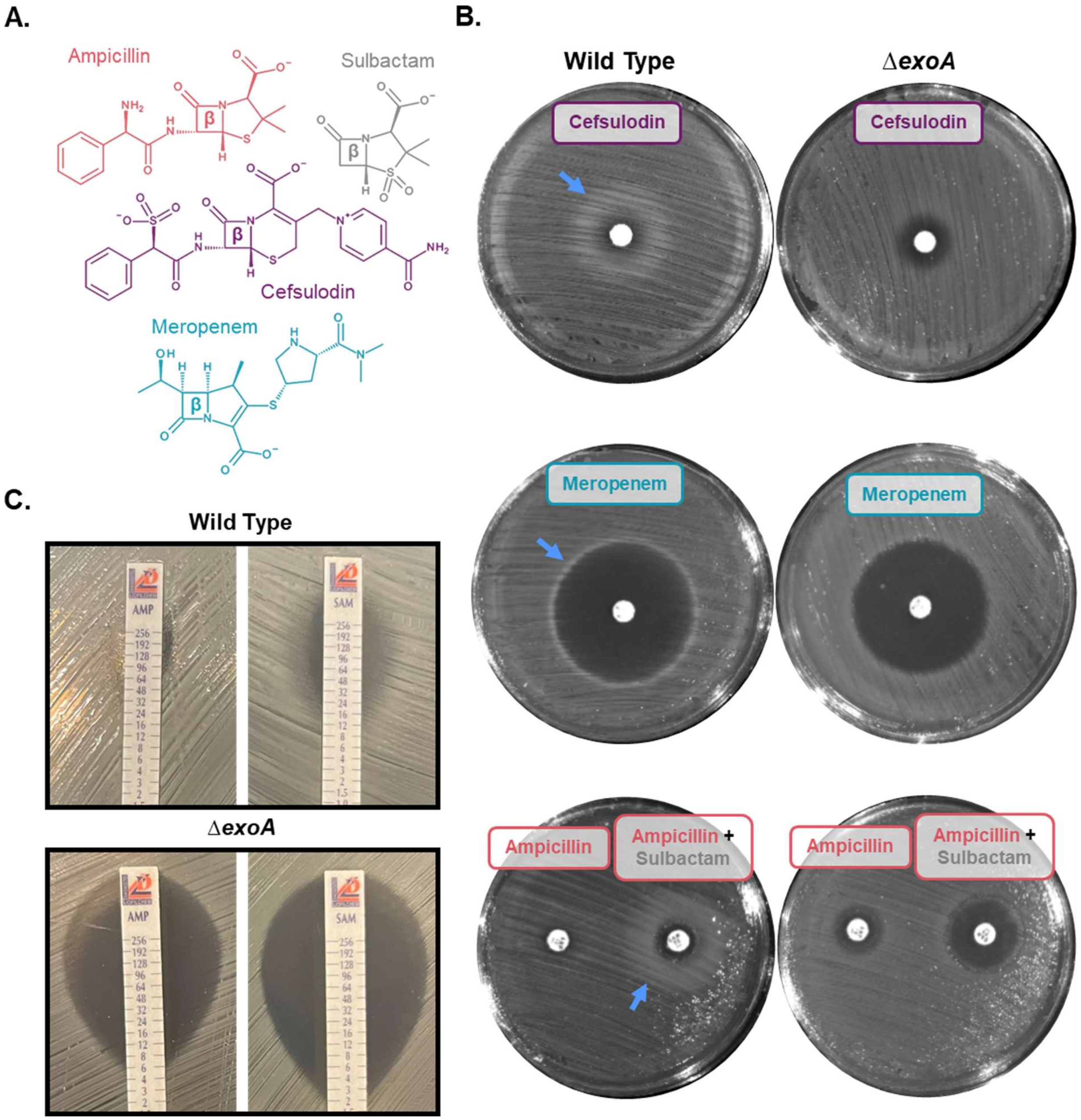


**Supplemental Figure 6. Succinoglycan is protective against β-lactam antibiotics.** A) Structure diagrams of ampicillin, sulbactam, cefsulodin, and meropenem are displayed with the β-lactam ring of each denoted with a β. B) ATGN media indicated strains applied on the surface prior to application of disks containing either 100 µg of cefsulodin (top), 10 µg of meropenem (middle), 10 µg ampicillin (bottom, left disk), or 10 µg ampicillin + 10 µg sulbactam (bottom, right disk). Blue arrow indicates ring of hazy growth due to the accumulation of succinoglycan as indicated by the absence in the *exoA* deletion strain. C) Minimum inhibitory concentration (MIC) determined by MIC strips of indicated strains to ampicillin (AMP, left) and ampicillin + sulbactam (SAM, right). Numbers on strips represent antibiotic concentration. Data for AMP is the same as shown in Figure 6.

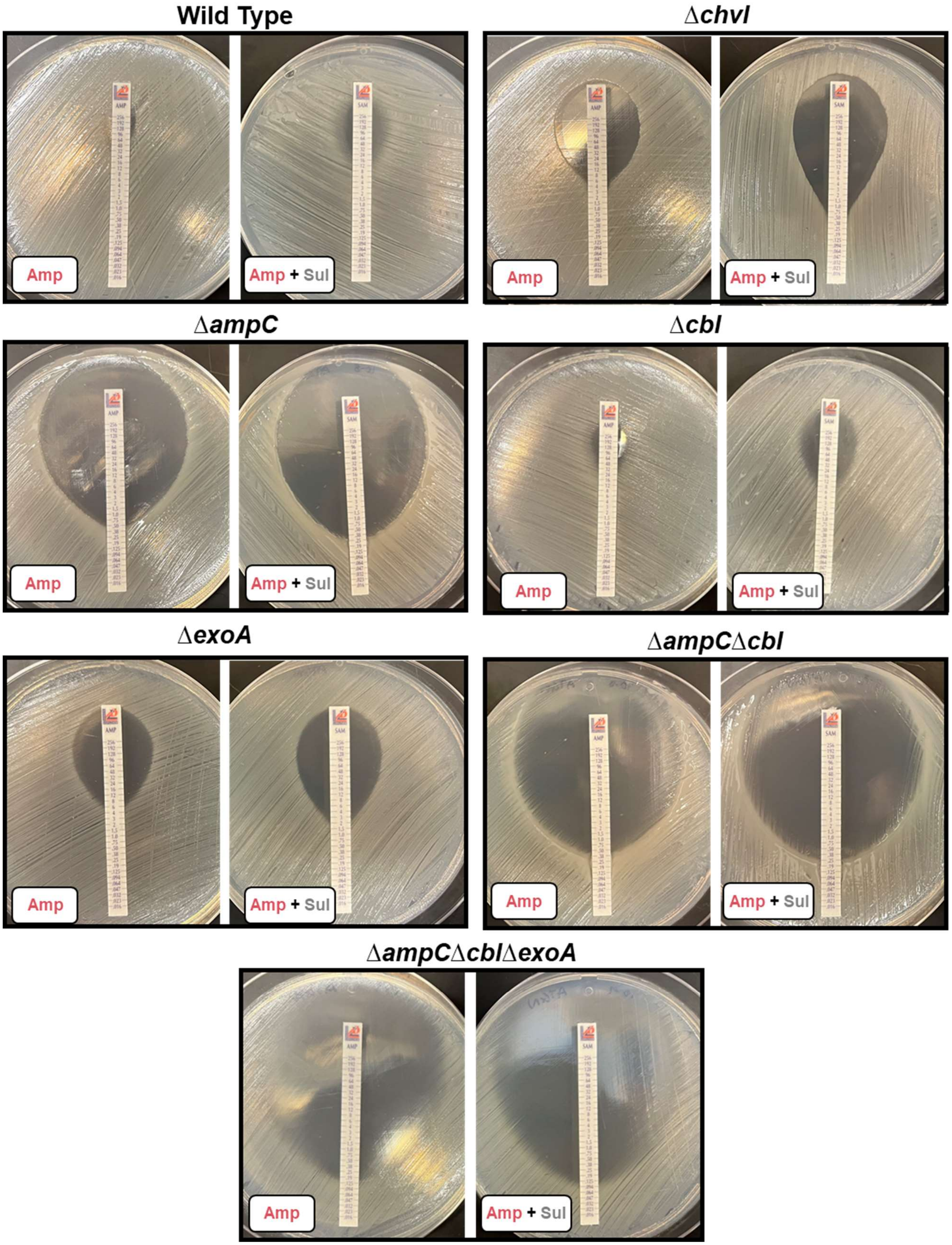
Wild Type
Amp
Amp + Sul
Δchvl
Amp
Amp + Sul
ΔampC
Amp
Amp + Sul
Δcbl
Amp
Amp + Sul
ΔexoA
Amp
Amp + Sul
ΔampCΔcbl
Amp
Amp + Sul
ΔampCΔcblΔexoA
Amp
Amp + Sul

**Supplemental Figure 7. AmpC and succinoglcyan confer resistance to ampicillin.** Minimum inhibitory concentration (MIC) determined by MIC strips of indicated strains to ampicillin and ampicillin + sulbactam. Numbers on strips represent antibiotic concentration. Some of the data for wildtype, ∆*exoA*, and ∆*cbl* are also shown in Figure 6, Supplemental Figure 6, and Figure 7.

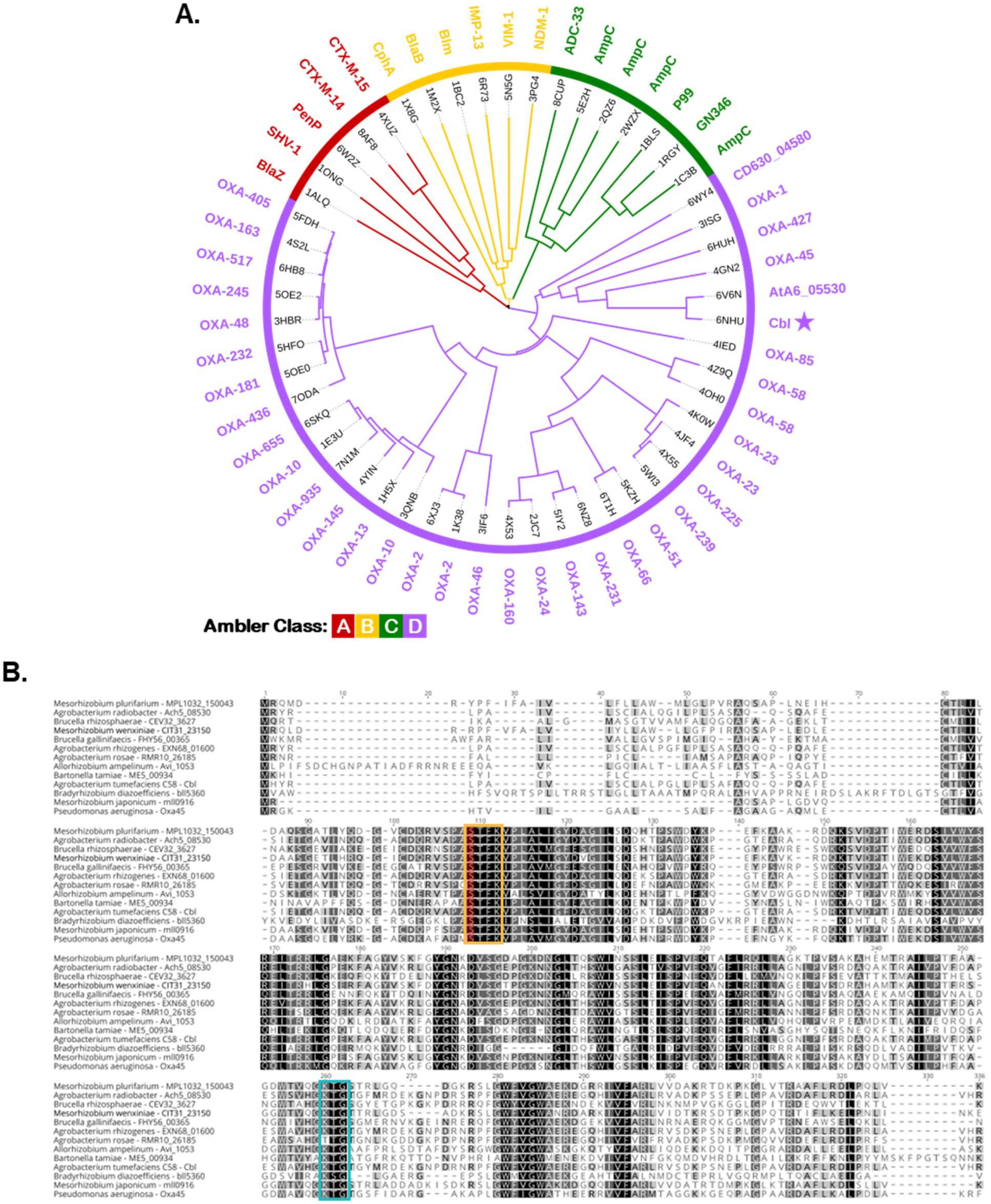
A.
B.
Ambler Class: A B C D
CTX-M-15
CTX-M-14
PenP
SHV-1
BlaZ
CphA
BlaB
Blm
IMP-13
VIM-1
NDM-1
ADC-33
AmpC
AmpC
AmpC
P99
GN346
AmpC
CD630_04580
OXA-1
OXA-427
OXA-45
AtA6_05530
Cbl
OXA-85
OXA-58
OXA-58
OXA-23
OXA-23
OXA-225
OXA-239
OXA-51
OXA-66
OXA-231
OXA-143
OXA-24
OXA-160
OXA-46
OXA-2
OXA-2
OXA-10
OXA-13
OXA-145
OXA-935
OXA-10
OXA-655
OXA-436
OXA-181
OXA-232
OXA-48
OXA-245
OXA-517
OXA-163
OXA-405
Mesorhizobium plurifarium - MPL1032_150043
Agrobacterium radiobacter - Ach5_08530
Brucella rhizosphaerae - CEV32_3627
Mesorhizobium wenxiniae - CIT31_23150
Brucella gallinifaecis - FHY56_00365
Agrobacterium rhizogenes - EXN68_01600
Agrobacterium rosae - RMR10_26185
Allorhizobium ampelinum - Avi_1053
Bartonella tamiae - ME5_00934
Agrobacterium tumefaciens C58 - Cbl
Bradyrhizobium diazoefficiens - bll5360
Mesorhizobium japonicum - mll0916
Pseudomonas aeruginosa - Oxa45

**Supplemental Figure 8. Cbl is a Class D β-lactamase conserved in some Hyphomicrobiales species. A)** Structural similarity dendrogram of crystal structures of β-lactamases of each Ambler class including Cbl,generated using Fold Tree (Moi *et al.*, 2023). Colors represent each class A-D. C) Amino acid sequence alignment of Cbl and orthologs from other species. Conserved Class D β-lactamase motifs are highlighted by the same colors as in panel A. The S60 active site is highlighted in red.

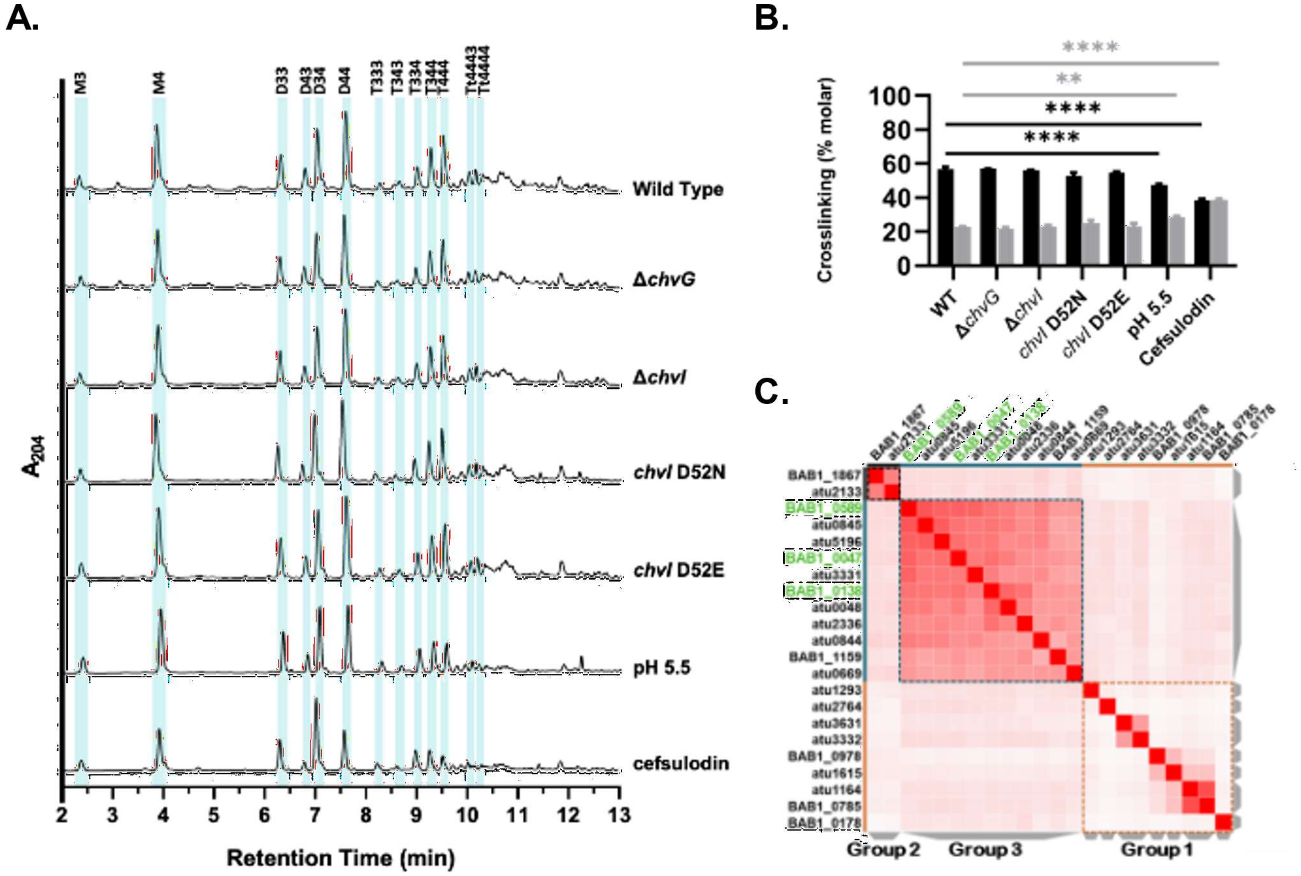


**Supplemental Figure 9. Muropeptide profiles reveal a lack of minimal modification to peptidoglycan by LDT enzymes.** A) HPLC traces of PG muropeptides from cells grown to late exponential phase (OD 0.5 – 0.8) in ATGN media. Strains include wild-type (WT), Δ*chvG*, Δ*chvI*, *chvI*$^{D52E}$, and *chvI*$^{D52N}$ cells. WT cells were also grown to late exponential phase in ATGN at a pH of 5.5 or in the presence of 25 µg/ml cefsulodin. B) Percent of peptidoglycan fragments containing an DD (black) or LD (gray) crosslink in late exponential to early stationary phase (OD 0.5 – 0.8) wild-type (WT), Δ*chvG*, Δ*chvI*, *chvI*$^{D52E}$, and *chvI*$^{D52N}$ cells. WT cells were also grown in ATGN at a pH of 5.5 or in the presence of 25 µg/ml cefsulodin to the same OD range for fragment analysis. A two-way ANOVA was used to compare differences in means of three replicates between each condition, followed by Šídák's multiple comparisons test with a single pooled variance. ns, not significant; *, p<0.1; **, p<0.01; ***, p<0.001; ****, p<0.0001. Only significant comparisons to WT are shown. Extended statistics can be found in supplemental table 14. C) Similarity matrix of LDTs from *Brucella abortus* and *A. tumefaciens*. Brucella LDTs highlighted in green are LDTs demonstrated to crosslink OMPs to PG in *Brucella abortus.* (Godessart *et al.*, 2020).
